# Assessing molecular, cellular and transcriptomic bases of laminar perfusion and cytoarchitecture coupling in the human cortex

**DOI:** 10.1101/2025.11.03.686411

**Authors:** Fanhua Guo, Chenyang Zhao, Ravi R. Bhatt, Zixuan Liu, Andy Jeesu Kim, Zidong Yang, Siyi Xu, Kay Jann, Mara Mather, Neda Jahanshad, Danny JJ Wang

## Abstract

Understanding how cellular architecture organizes cortical function requires mesoscopic approaches that resolve structure-function coupling in vivo. Here we introduce cerebral blood flow (CBF) and cell-body staining intensity (CSI) similarity index (CCSI), a localized similarity index between the CBF estimates from ultra-high-field 7T arterial spin labeling images and CSI from the BigBrain histology images to serve as a quantitative marker of laminar perfusion-cytoarchitecture coupling. CCSI revealed a reproducible, region-specific alignment between laminar vascular and cellular profiles across the cortex. Going beyond CBF, CCSI selectively tracked mitochondrial respiratory capacity and colocalized with capillary endothelial and mature non-myelination oligodendrocyte populations forming neurovascular interfaces. Transcriptomic enrichment highlighted pathways related to vascular remodeling, oxidative metabolism, and lipid-myelin homeostasis, indicating that CCSI reflects integrated metabolic-structural specialization. At the systems level, CCSI strengthened structure-function gradient correspondence in transmodal cortices, such as the default mode network. Together, these findings establish CCSI as a physiologically grounded, non-invasive marker of perfusion-cytoarchitecture alignment, providing a cross-scale framework linking cortical microstructure, metabolism, and functional organization.

## Introduction

The human brain is a dynamic system of distributed yet modular networks^1^, where structural connectivity provides the scaffold for functional interactions^2,3^. Advances in diffusion magnetic resonance imaging (MRI) tractography^4^ and functional MRI^5^ (fMRI), combined with graph-theoretic tools^6^, have revealed fundamental principles of macroscale (large-scale networks and interregional connectivity) organization^2^. However, recent studies have revealed a striking decoupling between macroscale structural and functional gradients in high-order networks^7,8^, indicating that structure alone cannot account for higher-order specialization. A key challenge is therefore to identify the additional organizational principles that bridge macroscale structural architecture with functional dynamics in the brain.

The past decades have seen major progress in understanding the human brain through microscale (cellular and subcellular components) technologies. High-resolution microscopy combined with AI-based analyses can reconstruct dense synaptic connections^9^ within small tissue volumes (1 mm^3^). Large-scale transcriptomic resources, such as the six-donor microarray dataset from the Allen Institute^10^, provide spatially resolved gene expression maps across the entire brain. Histological advances have yielded the BigBrain atlas^11^, an ultra-high-resolution (20 μm) 3D model capturing cytoarchitectonic features by silver-stained to highlight cell bodies. In parallel, metabolic mapping, including a human brain atlas of mitochondrial respiratory capacity and diversity^12^, has revealed regional variation in cellular energy supply. Together, these resources highlight the unprecedented detail now available at the micro-scale. Such molecular, cellular, and cytoarchitectonic information have yet to be translated into an understanding of the functional specialization and structural organization of brain networks, underscoring the need for mesoscale (laminar and columnar organization bridging cellular and network levels) information to bridge micro- and macro-levels.

The development of ultra-high-field (UHF) MRI was driven by the goal of imaging the brain at the mesoscale (sub-mm), bridging the gap between microscale synapses and neurons and macroscale regions and networks^13^. Recent advancements with UHF fMRI with blood-oxygen-level-dependent (BOLD) and vascular space occupancy (VASO) contrasts now enables the study of human brain function at the level of cortical layers and columns^14,15^. Yet the following challenges still remain in linking mesoscopic fMRI to neuronal activity: (1) the BOLD signal reflects complex interactions among cerebral blood flow (CBF), blood volume (CBV), and oxygen metabolism, limiting quantitative interpretation across layers with different baseline physiology^16^; (2) BOLD is highly susceptible to venous contamination, especially from pial veins on the cortical surface^14,16^; and (3) VASO-based CBV fMRI improves specificity but measures only relative CBV changes, confounded by layer- or column-dependent baseline CBV differences^16^.

CBF measured by arterial spin labeling (ASL) is a key in vivo marker of neurovascular function, as labeled arterial water exchanges with tissue water in capillaries near sites of neural activation^17^. Animal optical imaging shows that CBF can be rapidly regulated at the capillary level (e.g., via pericytes)^18,19^. ASL perfusion fMRI is able to visualize orientation columns in the cat visual cortex with superior spatial resolution to BOLD^20^, and its hemodynamic response precedes that of the BOLD signal by approximately one full second^21^. At 7T, ASL benefits from increased SNR (B0^1.65^) and longer tracer half-life, yielding a nearly fivefold signal to noise (SNR) gain over 3T^22,23^. Pilot laminar ASL at 7T has revealed higher resting perfusion in the middle cortical layers^24,25^, consistent with capillary density distributions derived from histology^26,27^. Because ASL is quantitative and originates from capillary/tissue compartments, high-resolution 7T ASL provides a mesoscale perfusion measure closely linked to neural activity.

In this study, we employed a novel turbo-FLASH (TFL) based rotated golden-angle stack-of-spirals (rGA-SoS) pseudocontinuous ASL (pCASL) sequence^28^ at 7T, enabling whole-brain 1-mm isotropic perfusion imaging, a spatial resolution and coverage that substantially surpass conventional ASL implementations^29^. Together with whole-brain high-resolution cell-body staining intensity (CSI) imaging from the BigBrain^11^ template, we proposed a new metric: the CBF-CSI similarity index (CCSI), which quantifies the coupling between laminar-specific CBF and cytoarchitectonic microarchitecture within local cortical regions (see Methods). Using this ASL sequence in 30 healthy participants, we mapped the distribution of both CBF and CCSI across the cortex. 14 out of 30 participants underwent retesting to verify the reproducibility of our results. We next examined how brain-wide heterogeneity in metabolic supply (CBF) and laminar perfusion-cytoarchitecture coupling (CCSI) related to microscopic maps, including mitochondrial complexes, spatial distributions of cell types, and diverse biological functions, thereby probing the physiological substrates of CCSI. Finally, we incorporated CBF and CCSI into structural and functional macroscale gradient analyses, demonstrating their potential to advance our understanding of how local microarchitecture and metabolism shape macro-scale brain function. Altogether, these findings highlight the potential of CCSI as a physiologically grounded biomarker for assessing laminar perfusion- cytoarchitecture coupling, with translational relevance for detecting alterations in vascular, metabolic, or glial organization and monitoring disease-related changes.

## Results

### Multiscale heterogeneity of CBF across cortical areas and lamina

We first estimated cerebral blood flow (CBF), defined as the volume of blood delivered to a unit of tissue per unit time, at the voxel level in the cortex for each participant using our newly developed rGA-SoS pCASL technique^28^. Regional mean CBF values within the cortical gray matter were then parcellated for each participant using segmentation tools in FreeSurfer and the HCP-MMP1.0 atlas^30^ (**Extended Data** Fig. 1a), yielding a 360 (regions) × 30 (participants) matrix. To minimize inter-individual differences in overall perfusion levels, CBF values were normalized (z-scored) within each participant. Principal component analysis (PCA) was then applied to this matrix, with the first principal component explaining 53.29% of the total variance. This component captures the dominant spatial pattern of CBF shared across participants (**Fig. 1a**). Among these regions, early sensory cortices (including primary visual, somatosensory, and auditory areas), the dorsolateral frontal cortex, posterior cingulate cortex, and the heavily myelinated area parieto-occipital sulcus area 2 exhibit high scores, whereas regions in the temporal lobe, such as the inferior temporal gyrus and portions of the middle temporal gyrus, show low scores. This spatial pattern aligns with observations reported in previous studies involving large cohorts using PET^31^ (**Extended Data** Fig. 2a) and low-res ASL-MRI^32^ (**Extended Data** Fig. 2b).

**Fig. 1.**
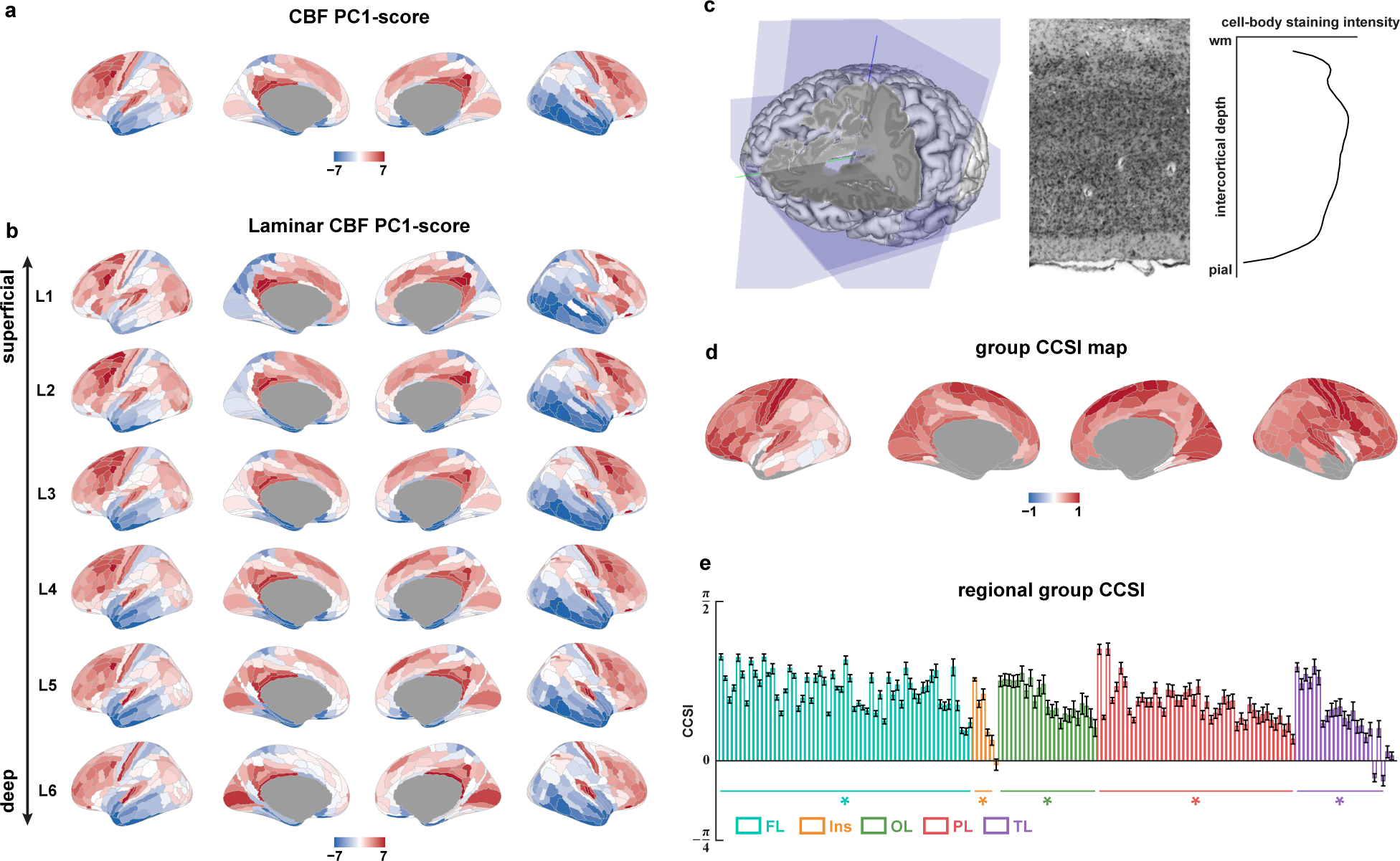
Spatial distribution heterogeneity of whole, laminar CBF-score and CCSI. **a**, Parcellation pattern of CBF principal component (PC1) scores across all participants (n = 30 independent biological replicates). **b**, Parcellation pattern of 6 laminar CBF PC1-scores across all participants. L1 to L6 are the six layers of gray matter from the superficial to the deep layer calculated by the equal volume algorithm. **c**, Schematic diagram of the distribution of laminar cell-body staining intensity (CSI) in brain histology of BigBrain data donors and the calculation of laminar profile. **d**, Average parcellation pattern of the CBF–CSI similarity index (CCSI) calculated from the laminar profiles of CBF and CSI across all participants after data quality control by SNR. **e**, Bar plot of CCSI distribution in each unilateral brain region across all participants (n = 30 independent biological replicates) after data quality control by SNR. Statistical significance was assessed using one-way RM-ANOVA and 2-side T-test (BH-FDR correction). The horizontal lines and asterisks below the bars indicate brain regions that are significantly diverge from 0 after multiple comparison correction (FDR < 0.05). Error bars indicate s.e.m. across participants. FL, frontal lobe; Ins, Insula; OL, occipital lobe; PL, parietal lobe; TL, temporal lobe.

We next segmented the cortical gray matter into 6 equi-volume layers (denoted as layers 1–6 from the superficial to the deep surface, **Extended Data** Fig. 1b) based on T1-weighted structural MRI data. This enabled the calculation of laminar CBF profiles across the 360 brain regions defined by the atlas. Using the same approach described above, we also computed the spatial distribution scores of CBF for each of the 6 cortical layers separately (**Fig. 1b**). The global CBF-score maps were broadly consistent across layers, yet a minority of regions showed clear layer-specific deviations. Among these regions, the spatial distribution score of the dorsolateral prefrontal cortex (dlPFC) and the dorsomedial prefrontal cortex (dmPFC) gradually decrease from the superficial to the deep layers. In contrast, early sensory cortices (including primary visual, somatosensory, and auditory areas) show a progressive increase in CBF-score from the superficial layer to deep layer.

Together, our results demonstrate that CBF exhibits multiscale heterogeneity across cortical areas and laminae, consistent with spatial variations in metabolic demand and functional specialization across the human cortex.

### Laminar-specific coupling between perfusion and cytoarchitectonic architecture

CBF is widely regarded as a physiological marker closely associated with regional energy supply^31^. Previous studies have shown that perfusion is strongly co-localized with the cytoarchitecture of the hippocampus, suggesting that CBF may be influenced by laminar microstructural features and potentially linked to local metabolic demand^33^. Based on this, we hypothesize that CBF is closely related to the distribution of cytoarchitectonic properties, particularly at the laminar scale. To investigate this relationship, we performed correlation analyses between laminar CBF profiles and cytoarchitectonic features from the BigBrain histological reconstruction (**Fig. 1c**). To ensure technical alignment with the CBF laminar profiles, we segmented the BigBrain cortical gray matter into 6 equi-volume layers (**Extended Data** Fig. 1c) and mapped it onto the same 360 brain regions using the same method described above. Given the limited number of laminar data points, we employed cosine similarity to quantify the resemblance between the laminar profiles of CBF and cell-body staining intensity (CSI) within each brain region for each participant. This measure, referred to as the CBF–CSI similarity index (CCSI, see Methods for calculation details), ranges from –π/2 to π/2, with positive values indicating aligned laminar profiles. Higher CCSI values reflect greater similarity between the laminar profiles of CBF and CSI.

After data quality control by SNR (**Extended Data** Fig. 3, see details in Methods), **Fig. 1d** presents the average bilateral CCSI map across participants and **Fig. 1e** presents the unilateral regional distribution of CCSI. Laminar profiles of CBF and CSI remained significantly correlated in nearly all cortical regions cross all participants (T-test with Benjamini-Hochberg false discovery rate correction; 59/59 in the frontal lobe, 5/6 in the insula, 23/23 in the occipital lobe, 46/46 in the parietal lobe, and 21/23 in the temporal lobe). Moreover, CCSI demonstrated significant region-specific heterogeneity (one-way repeated-measures ANOVA: main effect of brain region, F(156, 4524) = 24.21, p = 5.25x10^-43^, partial η^2^ = 0.46, Greenhouse-Geisser corrected). The group-level CCSI map exhibited broadly positive values across most cortical regions, with the strongest similarity along the central sulcus in primary sensorimotor areas and in occipital visual cortex. Moderate positive values extended into widespread association cortices, while lower or negative CCSI values were restricted to circumscribed regions of the inferior temporal and occipitotemporal cortex. **Extended Data** Fig. 4 shows the results based on the retest data of the 14 participants with two measurement sessions, which demonstrates the reliability and repeatability of CCSI and CBF-score maps obtained by our method.

Overall, CBF and CSI exhibit a high degree of similarity along the laminar depth direction, with substantial regional variation. Conceptually, CCSI may serve as a quantitative marker of the coupling between laminar scale cytoarchitecture and perfusion (i.e., blood flow and energy supply). These findings suggest that CCSI (a mesoscopic anatomic indicator) captures biologically meaningful laminar coupling between cytoarchitecture and perfusion. To probe the cellular and molecular substrates of this coupling, we next examined molecular, transcriptomic, and cell-type (micro-scale) correlates of CBF and CCSI maps.

### Laminar perfusion-cytoarchitecture coupling reflects mitochondrial metabolic capacity

Consistent with previous findings, the spatial distribution of CBF was highly correlated with cerebral metabolic rates of oxygen and glucose (**Extended Data** Fig. 2c,d), supporting its role as a proxy for local energy supply and metabolic activity^31^. Mitochondrial respiratory chain complexes, the core machinery of oxidative phosphorylation, are regionally heterogeneous across the human cortex^12^. To assess whether regional perfusion patterns and their laminar coupling with cytoarchitecture reflect underlying metabolic enzyme activity, we examined the spatial correspondence between CBF-score and CCSI maps and cortical distributions of mitochondrial Complexes I (NADH dehydrogenase, CI), II (succinate dehydrogenase, CII), and IV (cytochrome c oxidase, CIV).

**Fig. 2a** shows the spatial distribution maps of mitochondrial Complexes I, II, and IV (MCs), derived from MitoBrainMap^12^, which constructed whole brain maps by voxelizing human brain tissue, quantifying mitochondrial enzyme activity, and projecting to the cortex using multimodal imaging models. While mitochondrial complexes exhibit similar global gradient patterns across the cortex, they display distinct regional distributions, particularly in the temporoparietal junction, visual cortex, and parietal lobule. Although both CI and CII initiate the electron transport chain, they differ in their metabolic coupling: CI depends primarily on NADH^34^ from the tricarboxylic acid cycle, whereas CII receives electrons from FADH2 generated during succinate oxidation^35^. This distinction may account for the greater inter-regional variability of CI (1.27 ± 0.15, mean ± std) relative to CII (1.25 ± 0.09). **Fig. 2b** and **2c** show the Pearson correlation results between the spatial distributions of mitochondrial complexes and both the CBF-score and CCSI. Surprisingly, the CBF-score showed no significant correlation with any of the mitochondrial complexes, suggesting that CBF is shaped by multiple interacting factors^36^ (including vascular architecture, neural activity, and vasoregulatory processes) with oxidative phosphorylation capacity being only one contributor. In contrast, CCSI exhibited a significant positive correlation with CI, but not with CII or CIV. As CCSI quantifies the laminar-scale coupling between perfusion and cytoarchitectonic structure, its selective association with CI (compared to CII, which is less dependent on the tricarboxylic acid cycle, and CIV, which acts at the terminal step of the electron transport chain) suggests that metabolic entry via CI is more tightly aligned with laminar perfusion-cytoarchitectonic coupling. **Fig. 2d** presents the cortical map of tissue respiratory capacity (TRC), computed by integrating the distributions of mitochondrial complexes, along with its correlation with CCSI. The significant positive correlation (r = 0.30, variogram-p = 0.034) suggests that laminar-scale perfusion-cytoarchitecture coupling is associated with TRC. **Extended Data** Fig. 5a,b and **Fig.2e** show the distributions of mitochondrial density (MitoD) and mitochondrial respiratory capacity (MRC), along with their respective correlations with CCSI. While MitoD showed no significant spatial correlation with CCSI, MRC was strongly correlated (r = 0.32, variogram-p = 0.009), indicating that CCSI co-localizes mitochondrial respiratory capacity rather than mitochondrial abundance. **Extended Data** Fig. 5c-e shows correlations between the CBF-score map and TRC, MitoD, and MRC, without significant correlation. Together, these findings suggest that laminar perfusion-cytoarchitecture coupling, as captured by CCSI, is selectively coupled to tissue- or mitochondrion-level oxidative capacity, particularly via CI, potentially through yet-to-be-identified regulatory mechanisms.

**Fig. 2.**
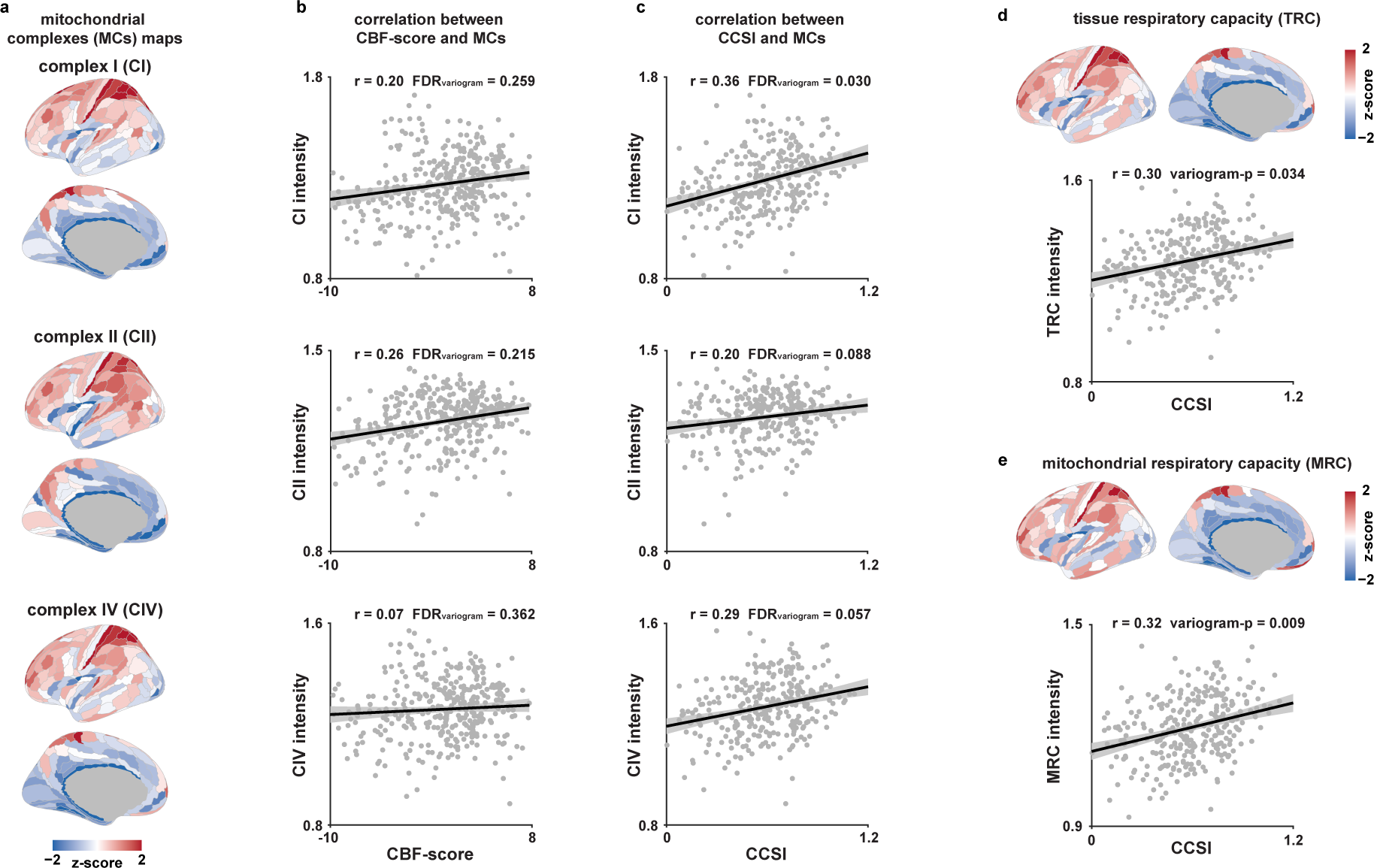
Spatial correlation analysis of CBF-score and CCSI map with mitochondrial complex and respiratory capacity related maps. **a**, Parcellation pattern maps of 3 mitochondrial complexes. These include complex I (CI, NADH–ubiquinone oxidoreductase), complex II (CII, succinate dehydrogenase), and complex IV (CIV, cytochrome c oxidase). **b**, Relationship between CBF-score distribution and mitochondrial complexes. We used Pearson correlation (r) and variogram-null model to calculate the p-value considering spatial autocorrelation. Exact r and FDR values after BH-FDR correction are marked in the figure. The shading is the 95% confidence interval and dots represent individual brain region (likewise in subsequent panels). **c**, Relationship between CCSI distribution and mitochondrial complexes. **d**, Parcellation pattern maps of tissue respiratory capacity (TRC) and relationship between CCSI and TRC. Exact r and variogram- null p value are marked in the figure. **e**, Parcellation pattern maps of mitochondrial respiratory capacity (MRC) and relationship between CCSI and TRC. Exact r and variogram-null p value are marked in the figure.

### Oligodendrocyte- and capillary-linked cellular architecture underlies laminar perfusion coupling

Distinct brain cell types exhibit different metabolic demands and energy burdens^37^, and the laminar coupling between perfusion and cytoarchitecture may reflect underlying cellular composition or function. Recent studies have leveraged microarray data from six donors of Allen Human Brain Atlas^10^ (AHBA) to infer the spatial distribution of major brain cell types and their associations with MRI-derived maps^38,39^. Here, we applied gene-category enrichment analysis^39^ (GCEA) to evaluate the spatial correspondence between our CBF-score and CCSI maps and the distribution of canonical cell types, and cross-validated the results using cell-class gene expression mapping^38^ (CGEM).

Based on the marker genes of cell-class in the brain by previous studies^40^, **Fig. 3a,b** presents the results of GCEA for the CBF-score and CCSI maps, while **Extended Data** Fig. 6a,b shows the corresponding results from CGEM. Both CBF and CCSI exhibited consistent and significant positive enrichments with endothelial cells (Endo). **Fig. 3c** further illustrates the spatial correspondence between the CGEM-estimated Endo-map and both perfusion-derived measures. Given that both CBF and CCSI are inherently vascular in nature, these results validate the sensitivity and specificity of the enrichment analyses in capturing cellular features. Notably, CCSI also demonstrated a robust and significant enrichment with oligodendrocytes (Oligo), whereas this relationship was absent for the CBF. **Fig. 3d** shows the spatial correspondence between the CGEM-derived Oligo-map and the CCSI distribution. This finding suggests that laminar-scale perfusion-cytoarchitecture coupling may be particularly linked to oligodendrocyte-related structure or function. Finally, CBF also showed a correlated distribution with inhibitory neuron subtypes In6a and In6b (**Extended Data** Fig. 6c,d), consistent with prior studies^41^. In6a and In6b cell types are fast-spiking parvalbumin-positive GABAergic neurons^42^ and this fast-spiking action is often considered to require higher energy support^43,44^.

**Fig. 3.**
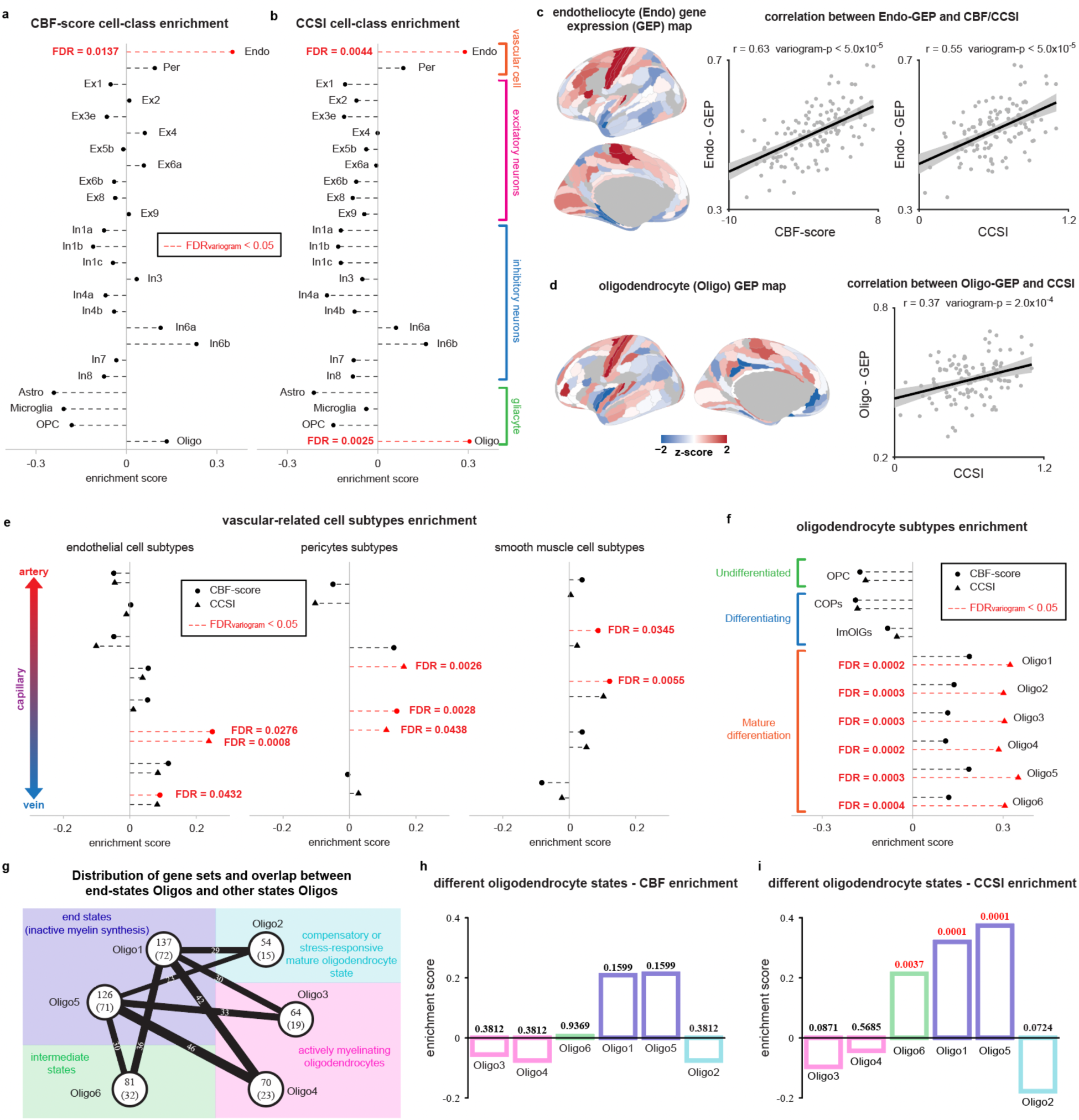
Cell-class enrichment analysis of CBF-score and CCSI. Whole cell-class enrichment analysis of CBF-score (**a**) and CCSI (**b**) by gene category enrichment analysis (GCEA). We used the variogram-null model to calculate the p value considering spatial autocorrelation. The exact FDR values that were significant after BH-FDR correction are marked in red (likewise in subsequent panels). Astro, Astrocytes; Oligo, Oligodendrocytes; OPC, Oligodendrocyte precursor cells; Endo, Endothelial cells; Per, Pericytes; Exa, Excitatory neuron subtype a; Ina, Inhibitory neuron subtype a. **c**, Spatial distribution of endothelial cells estimated using gene expression mapping and its relationship with CBF-score and CCSI. Exact r and p values calculated by the variogram-null model are marked in the figure. The shading represents the 95% confidence interval and dots represent individual brain region (same with **d**). **d**, Spatial distribution of Oligodendrocyte estimated using gene expression mapping and its relationship with CCSI. **e**, Vascular- related subtype cell-class enrichment analysis of CBF-score and CCSI by GCEA. These include various subtypes of three types of vascular cells: endothelial, pericyte, and smooth muscle cells. The vascular cells at the top come from arteries, the ones at the bottom come from veins, and the y-axis position of the capillaries in the middle also reflects the distance of each capillary cell from the artery or vein. **f**, Oligodendrocyte’s subtype cell-class enrichment analysis of CBF-score and CCSI by GCEA. COPs, Committed oligodendrocyte precursors; ImOlGs, Immature oligodendrocytes. **g**, Distribution of gene sets and overlap between end-states oligodendrocytes and other states oligodendrocytes. Each circle represents an oligodendrocyte in a certain state. The number in the circle indicates the number of genes in the marker gene set of that state. The number in the brackets indicates the number of genes remaining in the gene set after removing the overlap genes. The colored shading is based on the report of Jakel et al.^46^, which summarizes these six states into four functional states, including actively myelinating state, intermediate state, end state and oxidative stress state. **h**, Oligodendrocyte’s subtype cell-class enrichment analysis of CBF-score by redefined state-specific gene sets in **g**. The bar colors are consistent with the state classification in **g**. Exact FDR values calculated by the variogram-null model and BH-FDR correction are marked on each bar (same with **i**). **i**, Oligodendrocyte’s subtype cell-class enrichment analysis of CCSI by redefined state-specific gene sets in **g**.

We next performed enrichment analysis targeting vascular-associated^45^ and oligodendrocyte^46^ subtypes previously defined in single-cell transcriptome studies. **Fig. 3e** and **Extended Data** Fig. 6e show the enrichment results for vascular subpopulations derived from both analytical frameworks. Both CBF and CCSI were consistently and significantly enriched in vascular-associated cell types, particularly capillary-related subtypes across endothelial and pericyte, both of which are principal constituents of the neurovascular unit^47^. CCSI was more enriched in endothelial cells and pericytes but not in smooth muscle cells, while CBF was enriched in all vascular-related cells. Given that ASL-based CBF measurements predominantly reflect capillary perfusion, these results further support the validity of the enrichment analysis. **Fig. 3f** and **Extended Data** Fig. 6f display the enrichment results for oligodendrocyte subtypes. CCSI showed robust and consistent enrichment across multiple mature oligodendrocyte populations, but not in precursor, developing, or differentiating subtypes. In contrast, no significant associations were observed for CBF. This finding suggests that mature differentiation oligodendrocytes may be closely associated with laminar-scale coupling between perfusion and cytoarchitectonic organization, and that this relationship is likely independent of the total amount of local blood flow.

However, as noted by Jakel et al.^46^, the six oligodendrocyte categories (Oligo1–6) represent distinct transcriptional states rather than discrete cell subtypes. **Extended Data** Fig. 7a illustrates these six oligodendrocyte states^46^, along with the expression profiles of their respective marker gene sets and the degree of gene overlap among them. Thus, the significant enrichment of CCSI across all six oligodendrocyte categories observed in Fig. 3f may reflect a spurious effect driven by shared genes, rather than true state-specific associations. To address this, we removed all overlapping genes to generate non-redundant, state-specific marker sets (**Extended Data** Fig. 7a) and repeated the enrichment analysis. **Extended Data** Fig. 7b and 7c show the results for CBF and CCSI, respectively. After removing shared genes, CBF was significantly enriched for compensatory or stress-responsive oligodendrocyte states (FDRvariogram = 0.0057), whereas CCSI showed selective enrichment for the terminal (end-state, inactive myelin synthesis, FDRvariogram = 0.0006), intermediate-states (FDRvariogram = 0.0009) and compensatory or stress-responsive states (FDRvariogram = 0.0316) oligodendrocyte population. CCSI showed stronger enrichment in end-state oligodendrocytes, whereas results for other states were less reliable due to the small size of their gene sets. To address this, we only removed overlaps with the end-state gene set and redefined state-specific gene sets (**Fig. 3g**). Subsequent analyses revealed that only CCSI remained significantly enriched in end-state (FDRvariogram = 0.0001) and intermediate-state (FDRvariogram = 0.0037) oligodendrocytes (**Fig. 3h,i**). These findings suggest that laminar perfusion–cytoarchitecture coupling captured by CCSI is particularly associated with end-state oligodendrocytes and those transitioning toward end-states.

### Gene ontology signatures of perfusion-cytoarchitecture coupling and metabolic specialization

To further interpret the spatial heterogeneity of CBF and CCSI, we next performed gene ontology (GO) enrichment analysis to identify associated biological processes using aforementioned GCEA method (details in Methods). GO enrichment analysis of the CBF-score map (**Fig. 4a**) identified eight significantly enriched biological process terms. The enrichment of CBF-score for GO items involved in *negative regulation of Wnt/BMP signaling*, *LIF-mediated pathways*, *ion transmembrane transport*, *intracellular trafficking*, and *matrix mineralization*. This suggests that regional perfusion is aligned with molecular programs underlying cellular differentiation and developmental regulation. In addition, compared with the class I Histone Deacetylase (HDAC, an in vivo marker of cortical epigenetic regulation and plasticity constraints) expression map^48^ (**Extended Data** Fig. 2e) and the adolescent cortical scaling developmental expansion map^49^ (**Extended Data** Fig. 2f) in previous studies, the CBF-score showed weak but significant correlation with them. Together, these findings indicate that CBF heterogeneity may be shaped by metabolic demands associated with developmental processes during youth or ongoing cortical plasticity.

**Fig. 4.**
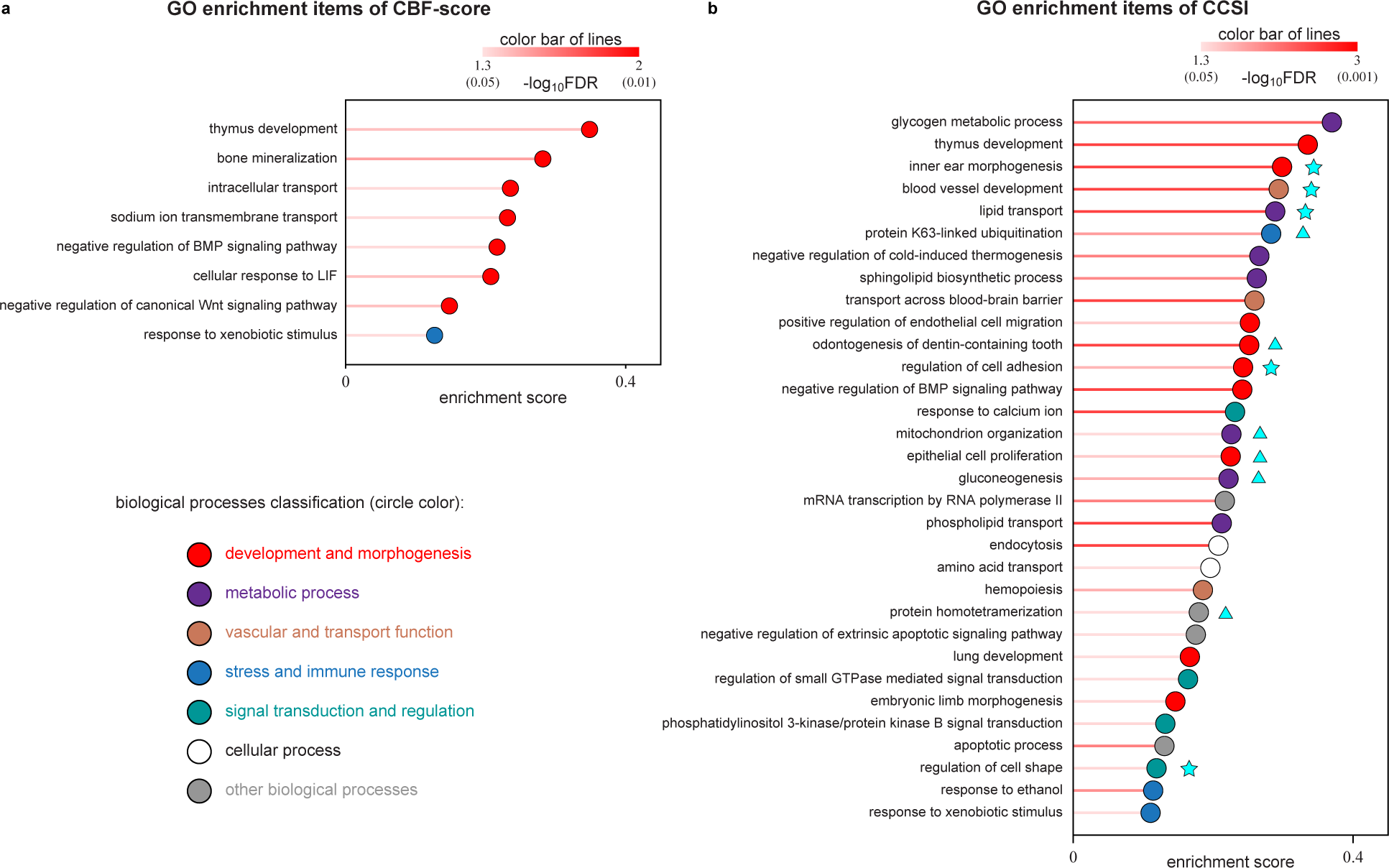
GO enrichment analysis of CBF-score and CCSI. **a**, CBF-score GO enrichment analysis of significant items. The figure shows the GO items that are significant after the variogram-null model and BH-FDR correction. The length of the line represents the enrichment score value, and the color of the line represents the enrichment FDR value. The color of the circles indicates the biological process classification to which these GO items belong (same with **b**). BMP, bone morphogenetic protein; LIF, leukemia inhibitory factor. **b**, CCSI GO enrichment analysis of significant items. A five-pointed star after a GO item indicates that the biological process of this GO item or the homologous signaling pathway expressed within this biological process is closely related to the neurovascular coupling process. A triangle after a GO item indicates that the biological process of this GO item or the homologous signaling pathway expressed within this biological process is closely related to improving mitochondrial metabolic capacity.

GO enrichment analysis of the CBF-score map (**Fig. 4b**) identified 32 significantly enriched biological process terms. These terms indicate that CCSI is closely associated with developmental, metabolic, vascular, and transport processes. Notably, biological processes represented by GO items such as *regulation of cell adhesion* and *blood vessel development* (highlighted by stars in **Fig. 4b**), or their homologous signaling pathways, are closely related to the morphogenesis of neurovascular coupling, suggesting that the laminar perfusion-cytoarchitecture coupling captured by CCSI may involve specific components of the neurovascular unit. Furthermore, biological processes represented by GO items such as *mitochondrion organization* (highlighted by triangles in **Fig. 4b**) are associated with enhanced mitochondrial metabolic capacity, suggesting potential signaling pathways that support the increased mitochondrial respiratory efficiency observed with CCSI.

### Reliability verification of the main findings

We examined the molecular, cellular, and transcriptomic correlates of CCSI and then assessed the reproducibility of our primary findings. Using the 14 participants’ retest data (**Fig. 5a–c**), we found that CCSI derived from the independent retest session also significantly colocalized with mitochondrial respiratory capacity and end-state oligodendrocyte distributions, confirming the reproducibility of our main results. Because our analysis assumes that the molecular (MitoBrainMap)^12^, cytoarchitectonic (BigBrain)^11^, and transcriptomic (AHBA)^10^ data from donors match the common distribution across multiple samples, we sought to determine the approximate sample size required to achieve robust and reliable results. To this end, we randomly selected subsets of 5, 10, 15, and 20 participants (20 iterations each) from the 30- participant dataset to replicate our primary findings. Robust and reproducible associations were consistently observed with sample sizes of ≥10 (**Fig. 5d-g**), underscoring the reliability and generalizability of our findings.

**Fig. 5.**
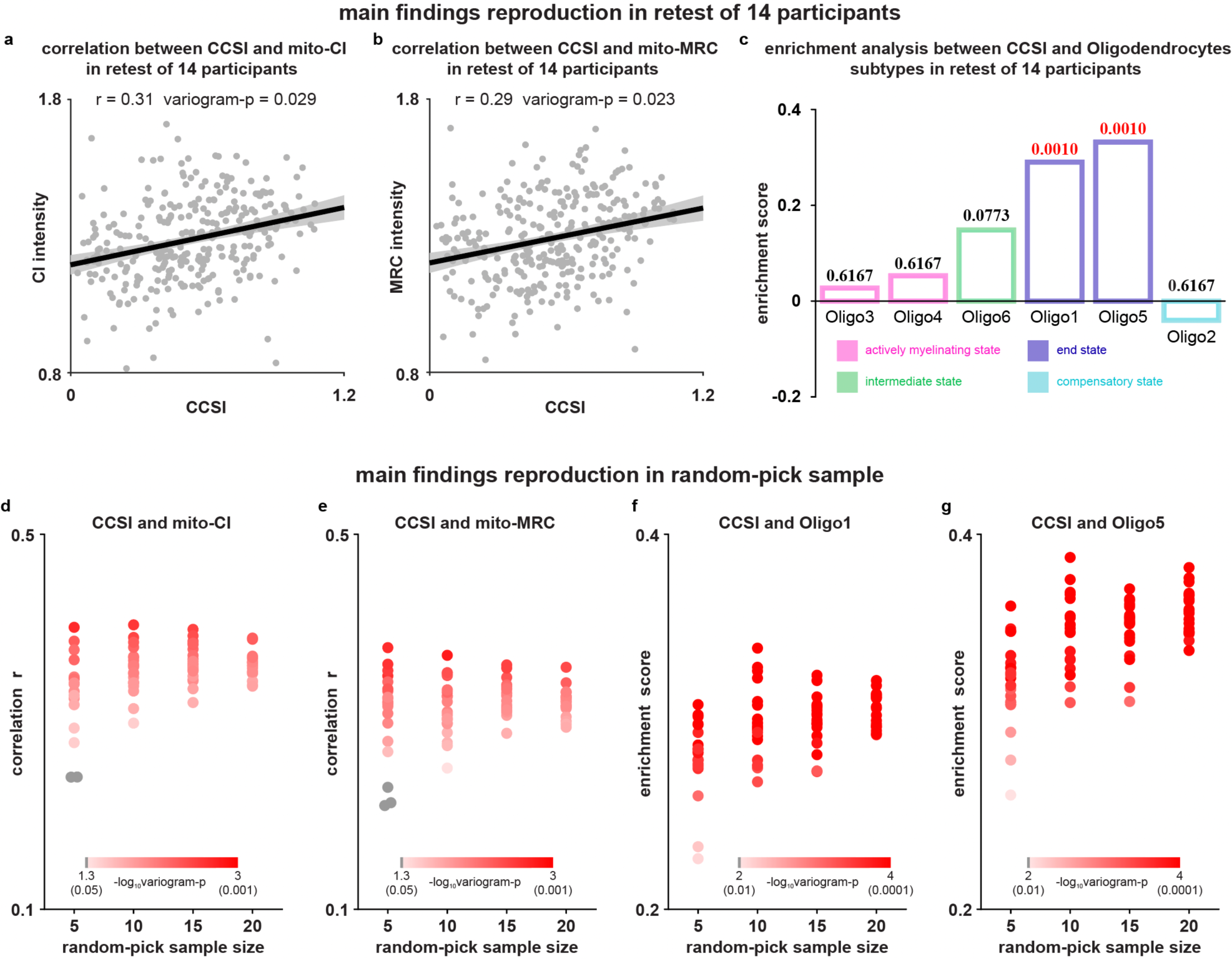
**Main findings are reproducible in the test-retest of 14 participants and the randomized samples. a-c**, Main findings are reproduced in the retest session of 14 participants. Relationship between CCSI distribution and mitochondrial CI (**a**) and MRC (**b**). We used Pearson correlation and variogram-null model to calculate the p-value considering spatial autocorrelation. Exact r and p values are marked in the figure. The shading is the 95% confidence interval and dots represent individual brain region. **c**, Reproduce the enrichment analysis results of CCSI and different oligodendrocyte states in Fig. 3i using 14 retest data. The bar colors are consistent with the state classification in Fig. 3g. Exact FDR values calculated by the variogram-null model and BH-FDR correction are marked on each bar. **d-g**, Main findings reproduction in randomly selected subsets of 5, 10, 15, and 20 participants (20 iterations each) from the 30-participant dataset. Each dot represents the main finding calculated after repeated sampling. The color of the dot represents the p-value calculated by the variogram-null model (corresponding to the color bar in the figure).

### CCSI strengthens structural-functional coupling in transmodal regions

We next examined whether laminar perfusion-cytoarchitecture coupling, as quantified by CCSI, contributes to understanding macro-scale patterns of cortical structure-function coupling. While studies have demonstrated a close correspondence between cortical microstructure and function^50,51^, recent human research has revealed a growing dissociation between structural and functional gradients in the transmodal networks^7,8^. To this end, we explored whether CCSI, a structural marker, and CBF, a metabolic measure, could help understand structure-functional coupling across the cortex.

**Fig. 6a** shows the parcellation map of the microstructure profile covariance gradient (GMPC) derived from BigBrain histological images^7^, and **Fig. 6b** presents the principal functional gradient^52^ (Gfunc) computed from resting-state fMRI data from the Human Connectome Project^53^ (HCP). **Fig. 6c** illustrates the spatial relationship between GMPC and Gfunc, as well as changes in model performance after adding CCSI as additional regressors and **Extended Data** Fig. 8 shows the changes in model performance after adding CBF as additional regressors. GMPC and Gfunc exhibit a moderate degree of spatial correspondence, consistent with prior studies^7^. Incorporating CBF as a co-regressor trended toward increased explained variance in Gfunc explained by the structural gradient (ΔR^2^, variogram-p = 0.055), suggesting that metabolic activity, as captured by CBF, may contribute to the functional embedding of cortical structure. While including CCSI did not enhance joint variance explanation, it might increase the contribution of GMPC to Gfunc prediction (ΔβG-MPC, variogram-p = 0.062), indicating that laminar perfusion-cytoarchitectural coupling may modulate the structural-functional relationship. Given that structural–functional coupling is known to degrade in the transmodal cortex, we separated GMPC into lower- and higher-order components (**Fig. 6d**) and repeated the analysis. As shown in **Fig. 6e,f**, structure-function coupling was much weaker in higher-order regions, consistent with previous reports^7,8^. In higher-order regions, CCSI (ΔβG-MPC, variogram-p = 0.021) significantly improved the predictive power of GMPC for Gfunc. These results suggest that CCSI may play a unique role in modulating structure-function relationships in the transmodal cortex, where traditional structural gradient predictors fall short.

**Fig. 6.**
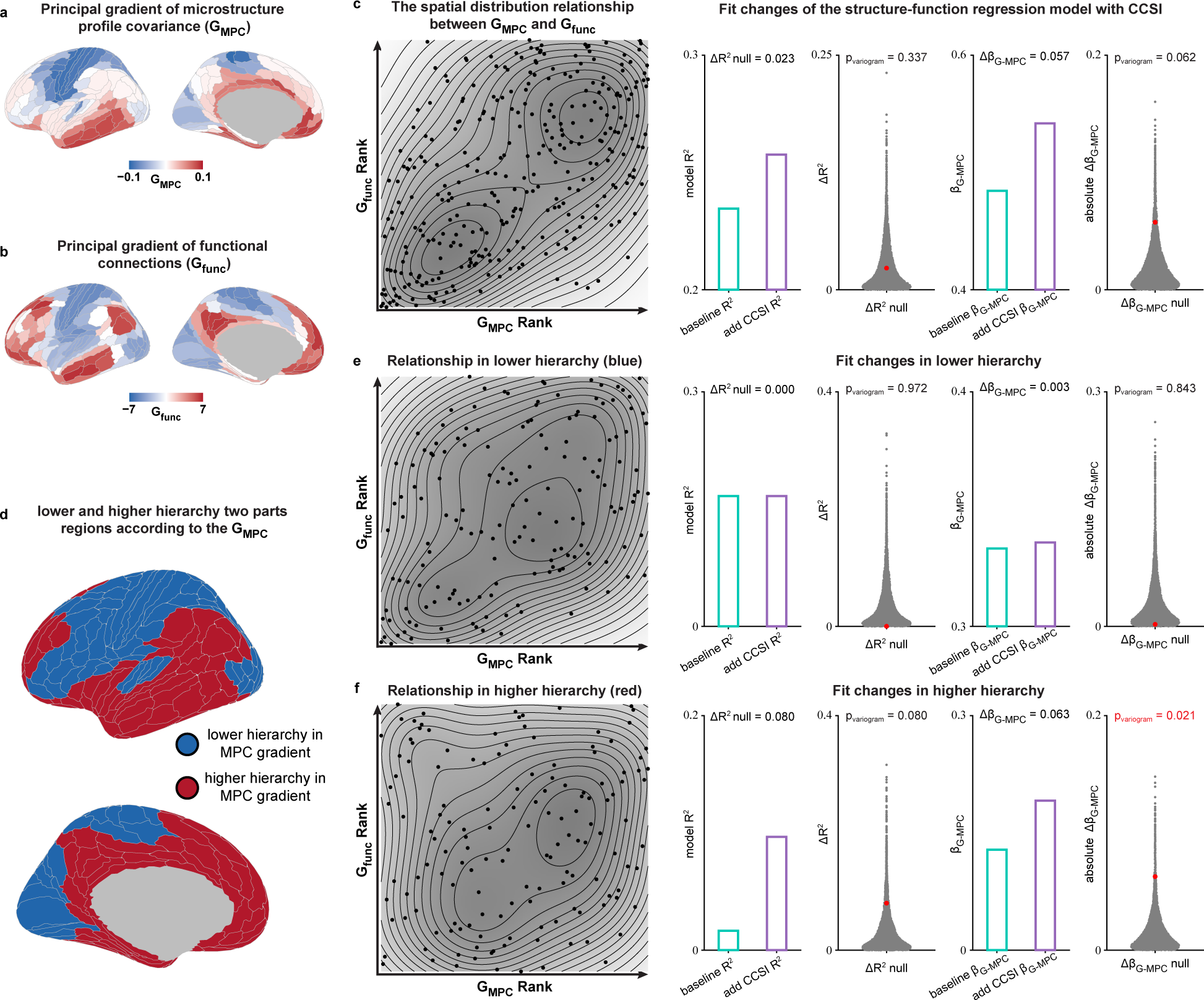
Fitting changes of the structure-function regression model with CCSI. **a**, Parcellation pattern map of principal gradient of microstructure profile covariance (GMPC). **b**, Parcellation pattern map of principal gradient of functional connections (Gfunc). **c**, The spatial distribution relationship between GMPC and Gfunc and the fit changes of the structure-function regression model with CCSI. Exact fitting parameters changes and p values calculated by the variogram-null model are marked in the figure. The cyan and purple bars represent the changes in the fitted parameters before and after CCSI was added as a regression parameter. The gray dots represent the changes in the fitted parameters distribution calculated using the variogram- null model, and the red dots represent the positions of the true values in the null distribution (same with **e** and **f**). **d**, Parcellation map of lower and higher hierarchy regions from GMPC. Blue regions are in the lower hierarchy and red regions are in the higher hierarchy. **e**, The spatial distribution relationship between GMPC and Gfunc and the fit changes of the structure-function regression model with CCSI in lower hierarchy. **f**, The spatial distribution relationship between GMPC and Gfunc and the fit changes of the structure-function regression model with CCSI in higher hierarchy.

## Discussion

Our study provides a multiscale characterization of cortical perfusion and its coupling with cytoarchitectonic structure, integrating ultra-high-field 7T ASL with histological, metabolic, and transcriptomic references. We demonstrate that cerebral blood flow (CBF) exhibits marked regional and laminar heterogeneity, reflecting spatial variations in metabolic demand and vascular organization. Beyond global perfusion patterns, we introduce the CBF-CSI similarity index (CCSI) as a quantitative and physiologically grounded marker of laminar perfusion-cytoarchitecture coupling. CCSI revealed robust region-specific alignment between perfusion and cellular density profiles, showing strong reproducibility across sessions and participants. Importantly, CCSI showed selective associations with mitochondrial respiratory capacity and with the spatial distributions of endothelial and mature oligodendrocyte populations that form neurovascular interfaces. Gene enrichment analysis further implicated pathways related to vascular remodeling, oxidative metabolism, and lipid–myelin homeostasis, suggesting that CCSI captures the integrated architecture of metabolic and structural specialization. At the systems level, CCSI enhanced structure–function correspondence in transmodal cortices, where conventional structural gradients show reduced explanatory power. Together, these findings establish CCSI as a noninvasive, mesoscopic biomarker that bridges vascular, cellular, and metabolic hierarchies, offering a new framework to investigate neurovascular–cytoarchitectonic alignment in the human cortex.

Our novel and reliable index of laminar perfusion–cytoarchitecture coupling (CCSI) addresses previous limitations in the sensitivity of conventional ASL to layer-specific vascular variation^29^. By integrating whole-brain high-resolution 7 T ASL with cytoarchitectonic profiles from BigBrain histology, we quantified laminar CBF profiles across the entire cortex and directly related them to the underlying cellular organization. This framework enables a quantitative evaluation of how local vascular supply aligns with microstructural architecture, offering new insights into the physiological principles governing neurovascular organization. In the following, we discuss the potential physiological mechanisms reflected by this metric.

Our CCSI primarily quantifies the extent of spatial correspondence between local vascular and neural cellular densities (including neurons and neurogliocytes). This metric reflects the spatially organized efficiency of the neurovascular unit (NVU), which provides the cellular and structural substrate for neurovascular coupling^54–57^ (NVC) and blood–brain barrier^56^ (BBB). The NVU is a multicellular and multi- scale dynamic network that spans distinct vascular segments and integrates neurons, astrocytes, oligodendrocytes, endothelial cells, pericytes, and smooth muscle cells into a coordinated signaling ensemble that facilitates the dynamic matching between neural activity and cerebral blood flow^55^. Regional variations in CCSI may therefore arise from differences in NVU compositions across cortical areas. Then, it raises the question of which NVU components play the dominant role in shaping the observed laminar perfusion-cytoarchitecture coupling.

Oligodendrocytes, derived from oligodendrocyte progenitor cells^58^ (OPC), are essential glial components that myelinate axons to accelerate conduction and reduce neuronal energy expenditure^58,59^. Beyond myelination, they sustain axonal metabolism by shuttling glycolytic substrates via monocarboxylate transporters^60–62^, buffer iron to mitigate oxidative stress^63^, and dynamically remodel myelin in response to neural activity^64,65^, thereby maintaining the structural and metabolic homeostasis of neural networks^59^.

Perineuronal oligodendrocytes (pn-OLGs) and perivascular oligodendrocytes (pv-OLGs) are specialized oligodendrocyte subtypes positioned in close anatomical proximity to neurons and blood vessels^66–69^, respectively. Early histological observations^66^ noted that pn-OLGs and pv-OLGs often coexist within the same microenvironment, forming structural bridges that link neuronal and vascular domains. Subsequent molecular and transcriptomic analyses revealed that the boundary between these two subtypes is indistinct, suggesting that they may represent spatially distinct manifestations or functional states within a common oligodendrocyte lineage^67^. Integrating molecular markers^67,68^ and gene-expression profiles^46,68^, both pn- and pv-OLGs are now recognized as belonging to the es-OLG class (mature, metabolically stable cells) whose spatial distribution shows strong correspondence with our CCSI map. During central nervous system development, OPC migrate along the vasculature under the guidance of endothelial CXCL12 signaling^70^ and subsequently differentiate into mature oligodendrocytes with pericyte support^71,72^. This developmental trajectory aligns with our enrichment findings showing that CCSI is significantly associated with endotheliocytes and pericytes gene expression at the capillary level. Transcriptome studies revealed that promoting intercellular adhesion is an important function of es-OLGs^46^. Recent connectomic reconstruction of the human cortex^9^ revealed that oligodendrocytes are the most abundant cell type and exhibit perivascular alignment across layers, supporting the notion that es-OLGs form widely distributed NVU structures linking vascular and neuronal domains. Within this framework, our results suggest that regions enriched in es-OLGs exhibit stronger laminar coupling between capillary distribution and cytoarchitectonic organization, as captured by CCSI. Taken together, these observations suggest that the regional heterogeneity of CCSI might primarily arise from NVUs composed of es-OLGs that physically couple neurons and capillaries, serving as key integrative hubs within the neurovascular network.

Although es-OLGs do not constitutively express high levels of myelin-synthesis genes, they retain the capacity to re-enter a myelinating state when required, suggesting a potential role in remyelination^46,67,68^. Rather than active myelin formation, their predominant functions involve sustaining signal transduction, mediating intercellular adhesion, and maintaining long-term cellular stability^46^. Through their dual associations with neurons and blood vessels, es-OLGs are well positioned to facilitate the bidirectional transfer of metabolic substrates, thereby supporting neuronal energy demands^67^. In parallel, their iron-buffering and antioxidative properties mitigate neuronal susceptibility to oxidative stress^46,63,73^. These metabolic and protective roles are consistent with the observed correlation between CCSI and mitochondrial respiratory capacity, and with the enrichment of CCSI in capillary-associated cellular populations. Perineuronal and perivascular OLG counts, or functional alterations have been reported across diverse neurological and psychiatric disorders^59,67–69^, including multiple sclerosis, schizophrenia, Alzheimer’s disease, and epilepsy. Together, these findings suggest that CCSI captures the spatial distribution of es-OLG-based neurovascular units and may serve as a potential imaging biomarker of neurovascular metabolic coupling in the human cortex.

CCSI provides a physiologically grounded index of laminar-scale perfusion–structure coupling with broad applicability. By leveraging high-resolution, non-invasive imaging, it enables investigation of cortical energy metabolism where histological validation is impractical. CCSI can sensitively probe regional heterogeneity in the alignment between vascular supply, cytoarchitectonic organization, and mitochondrial capacity. Because it reflects both vascular and oligodendrocyte-related processes, CCSI may be particularly informative for disorders such as multiple sclerosis, Alzheimer’s disease, and schizophrenia, where perfusion–structure coupling is disrupted. At the systems level, it offers a complementary metric linking structural gradients, metabolic demand, and functional connectivity. As a quantitative and physiologically interpretable measure, CCSI holds translational promise as a biomarker for detecting microvascular or metabolic alterations, monitoring disease progression, and assessing neurovascular or oligodendrocyte-targeted interventions.

Several limitations should be acknowledged. First, while the laminar resolution of in-vivo perfusion MRI remains limited, the derived CCSI maps were stabilized through multi-voxel depth averaging and cross-validation across participants. Second, CCSI was derived from in vivo perfusion data but referenced to a single postmortem BigBrain dataset, which may not reflect inter-individual variability in laminar architecture. However, by averaging across multiple participants, the group-level CCSI maps likely approximate population-level laminar patterns, mitigating potential biases from individual variability. Third, analyses of mitochondrial and cellular correlates relied on transcriptomic and histological atlases based on limited donor samples. Although cross-validation improved robustness, broader datasets are needed to confirm generalizability. Finally, as the present findings are correlational, future work integrating high-resolution diffusion MRI (e.g., 0.76 mm Connectome dMRI^74^, SANDI^75^) could generate individualized laminar maps and personalized CCSI indices, enhancing sensitivity to inter-individual variability and facilitating translational applications to behavior and disease.

In summary, we introduce the CBF-CSI similarity index (CCSI) as a physiologically grounded metric of laminar perfusion-cytoarchitecture coupling in the living human brain. Integrating high-resolution 7T ASL with cytoarchitectonic, metabolic, and cellular references, CCSI reveals meaningful associations with mitochondrial activity, oligodendrocyte organization, and structural-functional coupling, extending beyond conventional perfusion metrics. Despite methodological limitations, CCSI offers a promising across-scale framework bridging vascular, metabolic, and cellular dimensions of cortical organization. Future mesoscopic multimodal imaging will further refine this framework and advance its translational relevance for neuroscience and neurological disease.

## Methods

### Participants

Thirty volunteers (mean age = 25.9 years, SD = 5.1 years, 18 females) participated in this study. All participants in this study were healthy without known neurological, psychiatric disorders or major systemic diseases and refrained from caffeine 3 hours before the scan. All participants provided written informed consents according to a protocol approved by the Institutional Review Board (IRB) of the University of Southern California and were paid for attendance. Head motion was minimized by placing cushions on top and two sides of head. Fourteen of the participants underwent repeated scans on 2 different days for test and re-test.

### MRI data acquisition

MRI data were acquired on the investigational pTx part of a 7T Terra system (Siemens Medical Systems, Erlangen, Germany) with an 8Tx/32Rx head coil (Nova Medical, Cambridge, MA, USA). Anatomical images were acquired with a T1-weighted MP2RAGE sequence (0.7-mm isotropic voxels, 224 sagittal slices, FOV = 224×224 mm, TR = 4500 ms, TE = 3.43 ms, TI1 = 1000 ms, flip angle = 4°, TI2 = 3200 ms, flip angle = 5°, bandwidth = 200 Hz/Px, slice partial Fourier = 6/8, GRAPPA = 3).

For the perfusion image, a whole-brain 3D pCASL imaging with 1 mm isotropic resolution was performed at 7T using the FLASH-based rotated golden-angle stack of spirals (rGA-SoS)^28^ sampling scheme with following imaging parameters: FOV = 256 × 256 × 144 mm^3^, matrix size = 256 × 256 × 144, TR = 7 s, echo spacing (ES) = 19.5 ms, TE = 1.72 ms, ADC bandwidth = 400k Hz, ADC duration = 10.5 ms, flip angle = 15°, and echo train length (ETL) = 72, acceleration factor (R) = 4. Each M0, label, and control acquisition was acquired over 12 TRs, which was divided into 3 dynamic frames and can be reconstructed into 3 individual images to avoid lengthy temporal footprint, resulting in an effective temporal resolution for each image of 28 sec. In each TR, 1 spiral interleave with variable density (Rspiral = 2) in the kxy plane was acquired across 72 kz samples (Rz = 2). GA rotation (137.5°) was applied to spiral interleaves across 3D spatial space and dynamic frames, enabling superior incoherence in 4D. pCASL labeling has a duration of 1500 ms, and post labeling delay was 1800 ms. To avoid the loss of labeling efficiency due to field inhomogeneity at 7T. An optimized parameter set (Optimization of pseudo-continuous arterial spin labeling at 7T with parallel transmission B1 shimming) and a phase adjustment according to a prescan at the beginning of each scan (optimization of pseudo-continuous arterial spin labeling using off-resonance compensation strategies at 7T) was used.

### ASL reconstruction

we used a novel approach for ASL by integrating compressed sensing (CS) reconstruction with motion-resolved self-navigation to minimize motion artifacts^76^. Dynamic CS reconstruction incorporating spatial regularization on total variation (TVr) and temporal regularization on finite difference (FDt) across dynamic frames is performed to exploit the sparsity in 4D enabled by the sequence, which is described by,

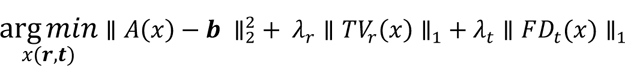

where x and b represent 4D image and k-space data, respectively. The forward operator A incorporates coil sensitivity weighting, non-uniform fast Fourier transform (NUFFT), and k-space undersampling. The inverse problem was solved using alternating direction method of multipliers (ADMM). Reconstruction was performed using 10 ADMM iterations with 3 nested conjugate gradient steps per iteration. Blurring caused by spiral sampling was mitigated by multi-frequency interpolation (MFI), and spiral trajectory errors were corrected by incorporating gradient impulse response (GIRF) that was measured on the scanner. Reconstruction was performed using 10 ADMM iterations with 3 nested conjugate gradient steps per iteration, requiring ∼90 secs and 14GB VRAM on a Nvidia RTX 5090 for each label/control image.

### MRI data preprocessing, parcellation and cortical layers definition

MRI data preprocessing were analyzed using AFNI^77^, FreeSurfer^78^, ANTs^79^, the mripy package^80^ (https://github.com/herrlich10/mripy) and MATLAB (R2021a). The preprocessing of the ASL data is following linear motion correction for control and label images, and registration average control image to T1w anatomical image. The perfusion image obtained by subtracting the label image from the control image can be used to obtain the CBF image according to the classic pCASL calculation method^81,82^. We estimated a 12-parameter linear transformation from the anatomical volume to a MNI space template by nonlinear transformation with ANTs. Subsequently, we used the HCP-MMP1 atlas^30^ in the MNI template space and transformed into the individual space of each participant using nonlinear transformation parameters to extract parcellation voxel-wise CBF signals by AFNI.

ASL images were upsampled to a finer grid of 0.5 mm^3^ isotropic spatial resolution to avoid singularities at the edges in angular voxel space using the ‘3dresample’ program in AFNI with the Nearest-Neighbor (NN) interpolation method. The T1w MP2RAGE anatomical volume was segmented into white matter (WM), gray matter (GM), and cerebrospinal fluid (CSF) using FreeSurfer’s automated procedure and its high-resolution option. A boundary-based algorithm in ANTs was used to co-register ASL control images to structural MP2RAGE images^83^. Six cortical layers in gray matter were calculated using AFNI/SUMA and custom python codes in mripy package with the equi-volume layering approach^84^. Although the cortical depths of different brain regions vary^85^ (From ∼4mm thickness of the prefrontal cortex to ∼2mm thickness of the primary sensory cortex) and the voxel size is not large enough to clearly distinguish the cortical layers, a large number of voxels randomly sampled at different cortical depth allowed us to derive a reliable continuous laminar profile of neural activity^25,80,86^. Finally, we can generate the laminar profile of CBF for 360 HCP-MMP1 brain regions in MATLAB.

### Histological data

We utilized the open-access BigBrain repository^11^, an ultra-high-resolution 3D reconstruction of a postmortem human brain stained for cell bodies, as the source of histological data. Their preprocessing pipeline included the following steps: First, the brain was paraffin-embedded and coronally sectioned into 7,400 slices, each 20 μm thick, which were then silver-stained to highlight cell bodies and digitized^87^. These images underwent manual inspection to identify artifacts such as rips, tears, shears, and stain crystallization. Subsequently, automatic correction procedures were applied, including nonlinear alignment to a postmortem MRI of the same individual acquired prior to sectioning, intensity normalization, and block averaging. Finally, the 3D reconstruction was achieved through a coarse-to-fine hierarchical procedure^11^. Computations were performed on inverted images, on which staining intensity reflects cellular density and soma size.

We downloaded the 3D volume at 200-μm resolution BigBrain data (https://bigbrain.loris.ca/main.php). Geometric meshes approximating the outer and inner cortical interface (i.e., the GM/CSF boundary and the GM/WM boundary) were generated for computing the depth^88^. Then we used the same aforementioned equi-volume algorithm to calculate the relative depth value of each gray matter voxel via the mripy package (https://github.com/herrlich10/mripy) based on the AFNI and FreeSurfer. Considering that the spatial resolution of our original ASL image is 1mm-isotropic, we also constructed 6 equivolumetric surfaces between the outer and inner cortical surfaces for BigBrain data. To map the HCP-MMP1 parcellations onto the BigBrain space, we used the BigBrainWarp toolbox^89^, which provides surface-based registration between BigBrain and standard MRI spaces. Subsequently, we generated the laminar profile of cell-body staining intensity (CSI) from Bigbrain data for 360 HCP-MMP1 brain regions.

### Laminar similarity calculation between CBF and CSI

To investigate the relationship between layer specific CBF and CSI, we conducted a correlation analysis of the laminar profiles of CBF and CSI across different ROIs. Layer 1 was excluded due to its low cellular density and the presence of large penetrating arteries, both of which may distort the CBF–CSI relationship. We employed vector cosine similarity to assess CBF–CSI relationship. Prior to calculating cosine similarity, we z-scored the five data points for both CBF and CSI profiles to eliminate baseline differences that could bias the vector angle. The similarity was then computed using the following formula:

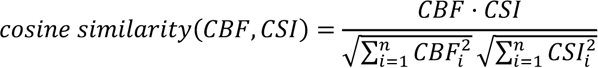

Here, CBF and CSI are 5-dimensional vectors representing cortical layers 2-6. Because cosine similarity values are bounded and skewed, Fisher z-transformation was applied prior to group-level statistical testing. For interpretability, we converted cosine similarity to angular distance using the arccosine transform and defined the CBF-CSI Similarity Index (CCSI) as:

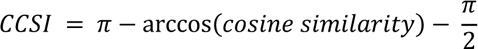

This transformation ensures that higher values indicate greater similarity. Specifically, we first inverted the arccos-derived similarity values (which range from 0 to π, with smaller values indicating higher similarity) by subtracting them from π, such that larger values represent greater similarity. We then centered the distribution by subtracting π/2, resulting in a final range of [–π/2, π/2], where 0 denotes no correlation and positive values indicate increasing similarity.

### CBF SNR estimation

To estimate the signal-to-noise ratio (SNR) of CBF images in a manner that accounts for both limited repetition and physiological variability, we employed a split-half method^90^ using interleaved repetitions. Specifically, we divided the CBF time series into odd and even subsets and calculated the mean CBF map for each. For each voxel, the noise standard deviation was estimated using the absolute difference between the odd and even means, scaled by their respective sample sizes:

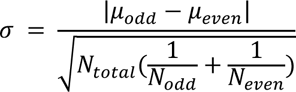

Where µodd and µeven denote the mean CBF images from odd and even repetitions, and Nodd, Neven are their respective counts. The median noise estimate across all voxels within each ROI was then taken as the representative noise level for that region. Finally, the regional SNR was computed as the ratio of the mean CBF value to the estimated noise standard deviation.

### CCSI quality control by SNR

**Extended Data** Fig. 3a presents the average bilateral CCSI map across participants. Overall, most cortical regions exhibit a high degree of similarity between CBF and CSI laminar profiles, with the exception of certain regions within the temporal lobe, where the similarity is relatively weaker. **Extended Data** Fig. 3b shows the distribution of group-level regional CCSI values after merging hemispheres. Except for the temporal lobe (30/37 regions), near all cortical areas showed significantly positive similarity after BH-FDR correction (67/67 in the frontal lobe, 5/6 in the insula, 24/24 in the occipital lobe, and 46/46 in the parietal lobe), broadly consistent with the CCSI map. Given the pronounced B0/B1^+^ inhomogeneities, increased susceptibility artifacts, and reduced coil sensitivity in inferior temporal regions at 7T^91^, the lower similarity observed in these areas may reflect diminished signal-to-noise ratio rather than true physiological differences. **Extended Data** Fig. 3c shows the average SNR map across participants. While the overall spatial distributions of CCSI and SNR differ across the cortex, low CCSI and low SNR co- localize in inferior temporal and orbitofrontal regions near the brain base. Across the entire cortical ROI, the correlation between CCSI and SNR was not statistically significant (Extended Data Fig. 3c, Pearson’s r = 0.2497, 2-side variogram-p = 0.2402). However, a significant positive correlation was observed in low-SNR regions (**Extended Data** Fig. 3d, SNR < 4, r = 0.3577, variogram-p = 0.0229), whereas no correlation was found in regions with higher SNR (**Extended Data** Fig. 3e, SNR > 4, r = -0.1054, variogram-p = 0.5004). Therefore, in the subsequent analysis of CCSI, we selected ROIs with SNR greater than 4 for analysis to exclude CCSI heterogeneity that may be caused by SNR.

### Mitochondria-related maps

We utilized recently published maps of mitochondrial respiratory capacity and diversity from Mosharov et al^12^, in which, a single postmortem brain from a neurotypical 54-year-old male donor was physically voxelized into 703 samples at MRI-comparable resolution (3×3×3 mm). Each voxel was biochemically profiled for mitochondrial density and oxidative phosphorylation capacity using assays of citrate synthase, mitochondrial DNA, and respiratory chain enzyme activities (Complex I, II, and IV), complemented by frozen tissue respirometry. These measures were integrated to generate indices of mitochondrial tissue density (MitoD), tissue respiratory capacity (TRC), and mitochondrial respiratory capacity (MRC = TRC/MitoD), thereby capturing both abundance and specialization of mitochondria across gray and white matter. The resulting voxel-wise mitochondrial features were registered to MNI stereotaxic space and upscaled to whole-brain maps using regression models linking biochemical measurements with multimodal MRI features. For the present study, we used the processed maps of CI, CII, CIV, MitoD, TRC, and MRC provided by the authors in MNI152 template space and resampled them to the HCP-MMP1.0 parcellation by AFNI to enable direct comparison with our imaging-derived metrics.

### Allen Human Brain Atlas (AHBA) gene expression data

We incorporated transcriptomic data from the Allen Human Brain Atlas^10^ (AHBA), using the optimized preprocessing pipeline^40^ described by Dear et al. Microarray measurements from six postmortem adult donors were first mapped to the HCP-MMP1 cortical parcellation (180 left-hemisphere parcels). To maximize cross-donor reproducibility, only parcels with samples from at least three donors were retained, yielding 137 cortical regions. Genes were ranked by differential stability (DS) across donors, and the top 50% (7973 genes) were selected to reduce noise. In the present study, we directly utilized the donor-consistent gene × region matrix, which represents a spatially reliable transcriptomic dataset. This matrix was used in its original form for integration with our imaging-derived measures.

### Variogram-null model generation

To assess the statistical significance of spatial correspondences while accounting for spatial autocorrelation, we generated null brain maps using the generative modeling framework proposed by Burt et al.^92^ and implemented in the neuromaps Python library^52^. This method constructs surrogate maps that preserve the empirical degree of spatial autocorrelation by matching the variogram of the observed data, while randomizing its topography. Specifically, the empirical map is first randomly permuted to destroy spatial structure, then spatial autocorrelation is reintroduced via distance-dependent kernel smoothing, with parameters optimized to minimize the error between empirical and surrogate variograms. Repeating this procedure yields a set of surrogate maps with spatial autocorrelation statistically indistinguishable from the empirical data but randomized spatial arrangement. Unless otherwise specified, we used the default settings and generated n = 20000 surrogates for each empirical map, which were subsequently used to derive null distributions of test statistics. To maintain the repeatability of our statistical estimates, we fixed the random seed at 1234. This approach provides spatially informed null models that offer more conservative and interpretable benchmarks than conventional permutation-based tests, which assume independence among regions.

### Gene category enrichment analysis (GCEA)

We performed gene category enrichment analysis (GCEA) following the ensemble-based framework proposed by Fulcher et al.^39,93^. For each gene, we calculated the spatial correlation between its AHBA-derived expression profile and the phenotype map of interest. Within each predefined gene set (GO and cell class), these correlations were Fisher’s z-transformed and summarized using the median as a robust category score, reducing sensitivity to a small number of highly deviant genes within each set. Statistical significance was assessed by comparing the observed category scores to null distributions as described in the variogram-null model generation section of the methods, with empirical p-values BH-FDR-corrected across categories.

### Cell-class gene expression mapping (CGEM)

When conducting cell category enrichment analysis, we also used the cell-class gene expression mapping^38^ (CGEM) method for cross-validation. For each cell category, a spatial gene-expression map was generated by computing the median expression value across all genes assigned to that category for each cortical region. The resulting category-level expression maps were then correlated with CBF or CCSI maps using Pearson’s correlation, and the correlation coefficients were Fisher z-transformed to yield enrichment scores. To assess statistical significance, null distributions of enrichment scores were constructed by repeating the same correlation using 20,000 variogram-null surrogate maps of CBF or CCSI that preserved spatial autocorrelation but randomized topography. P values were computed as the proportion of null scores exceeding the empirical enrichment score for each cell class.

### Gene set selection of GO items

To derive functional gene sets for enrichment analysis, we mapped the 7,973 reliably expressed genes retained from the AHBA preprocessing to Gene Ontology^94^ (GO) annotations using the R package *biomaRt* interfacing with the Ensembl database^95^. Gene symbols were first standardized (e.g., resolving ambiguous aliases such as MARCH and SEPT families) before querying. For each gene, associated GO identifiers, term names, and namespaces were retrieved via the *getBM* function. Only terms belonging to the Biological Process ontology were retained, while Molecular Function and Cellular Component categories were excluded. To ensure statistical robustness, we further restricted the analysis to GO categories containing at least 20 AHBA-mapped genes. The resulting curated collection of GO gene sets was used for downstream enrichment analyses.

### Gene set selection of cell class

Gene sets for major cortical cell types were obtained from Lake et al.^42^, who profiled over 60,000 single nuclei from human cortex and identified cell-type-specific marker genes distinguishing excitatory and inhibitory neurons, astrocytes, oligodendrocytes, OPCs, microglia, and endothelial cells. Vascular-associated cell subtypes, including endothelial and mural (pericyte and smooth-muscle), were derived from Garcia et al.^45^. Oligodendrocyte lineage subclusters, including mature and end-state oligodendrocytes, were obtained from Jäkel et al.^46^.

Because the gene sets representing different mature oligodendrocyte states showed substantial overlap, direct enrichment analyses risked confounding effects driven by shared genes rather than state-specific expression. To enhance the specificity of oligodendrocyte subtype comparisons, we implemented a two-step gene-set refinement procedure. First, we removed duplicate genes shared between any two of the six mature oligodendrocyte state gene sets to obtain non-overlapping subtype-specific gene lists, as shown in **Extended Data** Fig. 7a. This stringent filtering substantially reduced gene-set sizes for certain subtypes, occasionally leading to unstable or unreliable enrichment estimates. Next, given that CCSI showed stronger associations with mature oligodendrocytes biased toward end-state phenotypes, we further refined specificity by removing genes shared between the end-state oligodendrocyte gene set and all other oligodendrocyte states, yielding the final gene-set distribution shown in **Fig. 3g**.

### Other cortical maps

The PET-CBF, ASL-CBF, CMRO2, and CMRglc maps used in **Extended Data** Fig. 2 were obtained from previously published datasets^31,32^ compiled within the neuromaps toolbox^52^. The functional gradient map shown in Fig. 6b was also derived from the neuromaps toolbox^52^. The cell microarchitecture and structural gradient map presented in Fig. 6a were generated from BigBrain cell body staining intensity^11^ profiles processed using the ENIGMA toolbox^96^.

### Structural-functional coupling analysis

To assess whether CCSI or CBF contributes to large-scale functional organization, we quantified the relationship between microstructural and functional gradients across cortical parcels of the HCP–MMP1.0 atlas. The first model estimated the baseline structure-function correspondence using a simple linear regression of the functional gradient (Gfunc) on the microstructural gradient (GMPC):

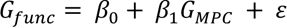

We then added either CBF or CCSI as an additional predictor in a multiple linear regression model:

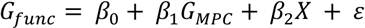

where X denotes the regional CBF or CCSI value. The improvement in model fit (ΔR^2^) quantified the additional variance in Gfunc explained by CBF or CCSI beyond microstructural gradients. Changes in the GMPC coefficient (Δβ1) were further examined to evaluate whether CBF or CCSI modulated the strength of structure-function coupling. Statistical significance of ΔR^2^ and Δβ1 was assessed using 20,000 permutations of variogram based null models.

### Statistical analysis

All statistical analyses were performed in MATLAB (R2021a). Group-level statistical significance of regional CCSI values was assessed using one-sample t-tests against zero, with p-values corrected for multiple comparisons using the Benjamini-Hochberg false discovery rate (BH-FDR). A one-way repeated-measures ANOVA was applied to evaluate region-specific differences in CCSI, with Greenhouse-Geisser correction applied where sphericity was violated. Prior to conducting the ANOVA test, each dataset was evaluated for normality using the Shapiro-Wilk test and for homogeneity of variance using the Levene’s test. Spatial correlations between imaging-derived maps (e.g., CBF, CCSI) and histological, metabolic, or transcriptomic references were evaluated using variogram-based permutation tests (n = 20,000) implemented in the neuromaps using the variogram framework proposed by Burt et al^92^, which preserves spatial autocorrelation of cortical topology. For gene and cell-class enrichment analyses, empirical correlations were Fisher z-transformed and compared against null distributions generated from the same variogram based null permutations, with significance determined after BH-FDR correction across categories. In the structure–function coupling analysis, model improvements (ΔR^2^) and coefficient changes (Δβ1) were evaluated via identical permutation tests. Unless otherwise specified, all tests were two-tailed with a significance threshold of p < 0.05 and corrected for multiple comparisons using the BH-FDR.

## Data Availability

The raw data of MP2RAGE and ASL used in this study can be obtained from OpenNeuro (https://openneuro.org/datasets/ds006871).

## Code Availability

The custom analysis code and the plotting code developed in this study is freely available at GitHub (https://github.com/FanhuaGuo/ASL-Bigbrain-CCSI.git).

## Acknowledgements

This work was supported by US National Institute of Health (NIH) grant UF1-NS100614, S10- OD025312, R01-NS114382, R01-EB032169, RF1AG084072, R01-EB028297, R01-NS134712 and R01- NS121040.

## Author Contributions

F.G. and D.J.W. conceived the study. F.G. and D.J.W. designed experiments. F.G., C.Z., Z.L. and A.J.K. executed experiments. F.G., C.Z., and R.R.B. analyzed experiments. C.Z. provided support to ASL. F.G. and D.J.W. drafted the manuscript. F.G., C.Z., R.R.B., Z.L., A.J.K., Z.Y., S.X., K.J., M.M., N.J. and D.J.W. interpreted the results, reviewed the paper, and approved the decision to submit the paper.

## Supplements

**Extended Data Fig. 1.**
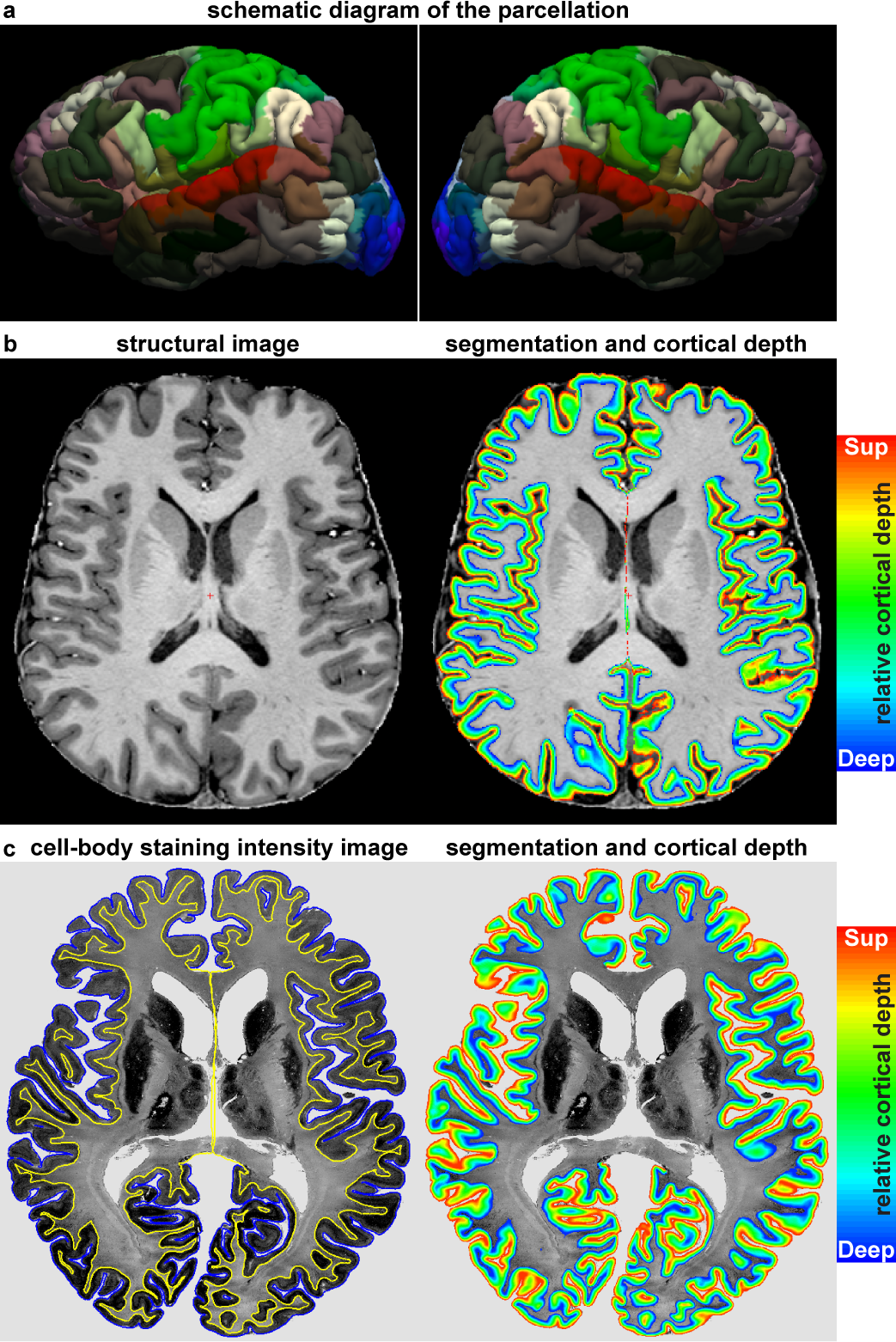
Schematic diagram of parcellation, segmentation and estimated cortical depth. **a**, Schematic diagram of parcellation by HCP-MMP1 atlas^30^. **b**, The left image is the anatomical image from T1w MP2RAGE sequence. The right image is the results of the segmentation and estimated cortical depth. **c**, Schematic diagram of cell-body staining intensity map^11^ and its calculated cortical depth.

**Extended Data Fig. 2.**
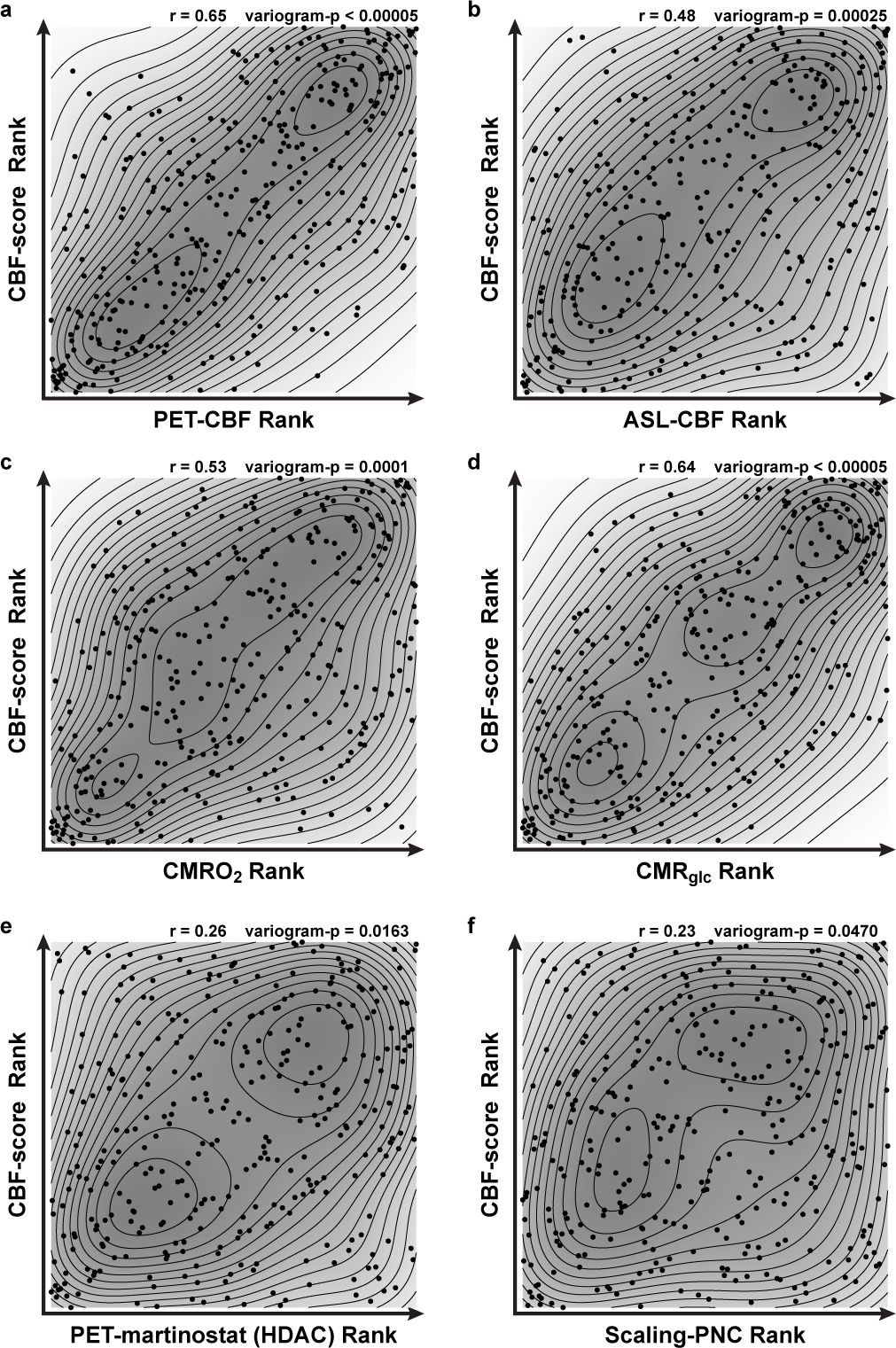
The spatial distribution relationship between CBF-score and PET-based CBF, ASL-based CBF, CMRO2 and CMRglc maps in previous studies. The spatial distribution relationship between CBF-score and PET-based CBF (**a**), ASL-based CBF (**b**), CMRO2 (**c**) and CMRglc (**d**) maps in previous studies. The points represent the values of each parcel, the black lines are equipotential lines, and the shaded gray values represent the probability density distribution. We used Pearson correlation and the variogram-null model to calculate the p-value considering spatial autocorrelation. Exact r and variogram-p values are marked in the figure (likewise in subsequent panels). **e**, The spatial distribution relationship between CBF-score and PET-martinostat. Martinostat PET maps measure class I Histone Deacetylase (HDAC) expression, an in vivo marker of cortical epigenetic regulation and plasticity constraints^48^. **f**, The spatial distribution relationship between CBF-score and scaling-PNC. Scaling-PNC quantifies region-wise surface area scaling relative to total cortical size in Philadelphia Neurodevelopmental Cohort (PNC) data in adolescents^49^.

**Extended Data Fig. 3.**
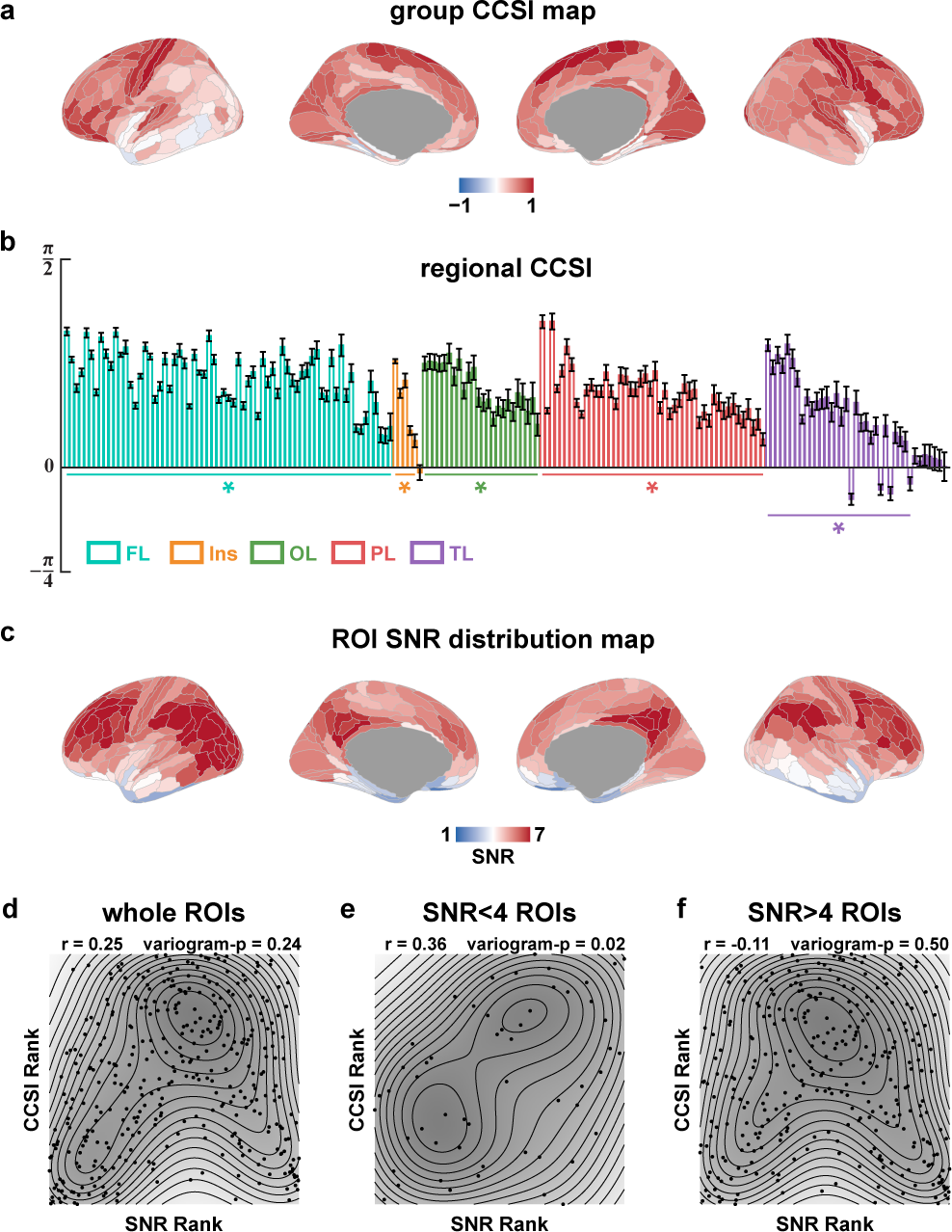
Spatial distribution pattern of CCSI and quality control based on SNR. **a**, Average parcellation pattern of the CBF–CSI similarity index (CCSI) calculated from the laminar profiles of CBF and CSI across all participants. **b**, Bar plot of CCSI distribution in each unilateral brain region across all participants (n = 30 independent biological replicates). Statistical significance was assessed using one-way RM-ANOVA and 2-side T-test (BH-FDR correction). The horizontal lines and asterisks below the bars indicate brain regions that are significantly greater than 0 after correction (FDR < 0.05). Error bars indicate s.e.m. across participants. **c**, Average parcellation pattern of the SNR across all participants. **d-f**, The spatial distribution relationship between CCSI and SNR in whole brain (**d**), low-SNR parcels (**e**), and high-SNR parcels (f). The points represent the values of each parcel, the black lines are equipotential lines, and the shaded gray values represent the probability density distribution. We used Pearson correlation and the variogram-null model to calculate the p-value considering spatial autocorrelation. Exact r and variogram -p values are marked in the figure.

**Extended Data Fig. 4.**
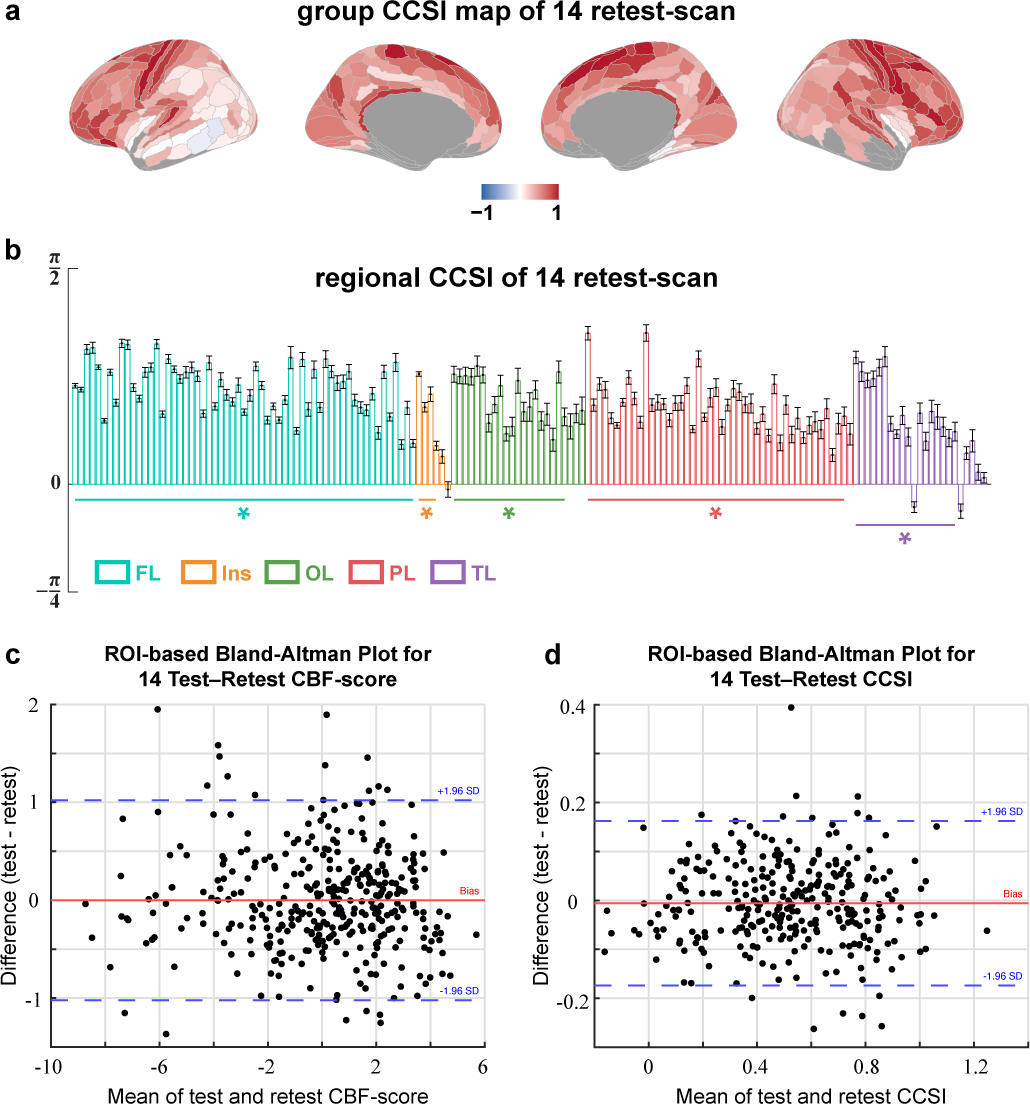
Retest reliability analysis of 14 participants. **a**, Average parcellation pattern of the CBF-CSI similarity index (CCSI) calculated from the laminar profiles of CBF and CSI across 14 participants retest session after data quality control by SNR. **b**, Bar plot of CCSI distribution in each unilateral brain region across 14 participants retest session (n = 14 independent biological replicates) after data quality control by SNR. Statistical significance was assessed using one-way RM-ANOVA and 2-side T-test (BH-FDR correction). The horizontal lines and asterisks below the bars indicate brain regions that are significantly greater than 0 after correction (FDR < 0.05). Error bars indicate s.e.m. across participants. Laminar profiles of CBF and CSI remained significantly correlated across nearly all cortical regions (59/59 in FL, 4/6 in Ins, 20/23 in OL, 45/46 in PL, and 18/23 in TL). Moreover, CCSI demonstrated significant region-specific heterogeneity (one-way repeated-measures ANOVA: main effect of brain region, F(156, 2028) = 14.02, p = 3.16x10^-14^, partial η^2^ = 0.52, Greenhouse-Geisser corrected). **c**, ROI-based Bland-Altman plot of CBF-score across 14 participants test and retest sessions. The red and blue lines are the bias between 2 sessions and the 95% confidence interval, and dots represent individual brain regions (same with **d**). **d**, ROI-based Bland-Altman plot of CCSI across 14 participants test and retest sessions.

**Extended Data Fig. 5.**
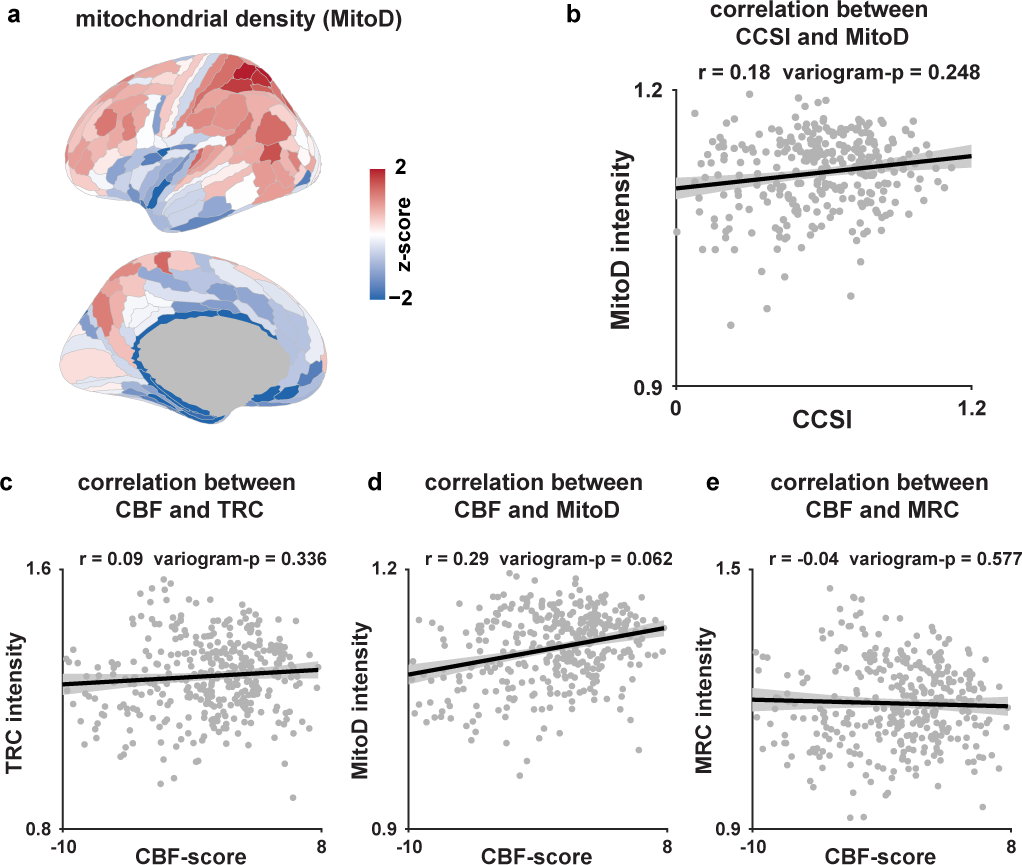
Mitochondrial density map and spatial correlation of CBF-score and CCSI map with mitochondrion-related maps. **a**, Parcellation pattern maps of mitochondrial density (MitoD). **b**, Relationship between CCSI distribution and MitoD. We used Pearson correlation and the variogram-null model to calculate the p-value considering spatial autocorrelation. Exact r and variogram-p values are marked in the figure. The shading is the 95% confidence interval and dots represent individual brain regions (likewise in subsequent panels). Relationship between CBF-score distribution and TRC (**c**), MitoD (**d**) and MRC (**e**).

**Extended Data Fig. 6.**
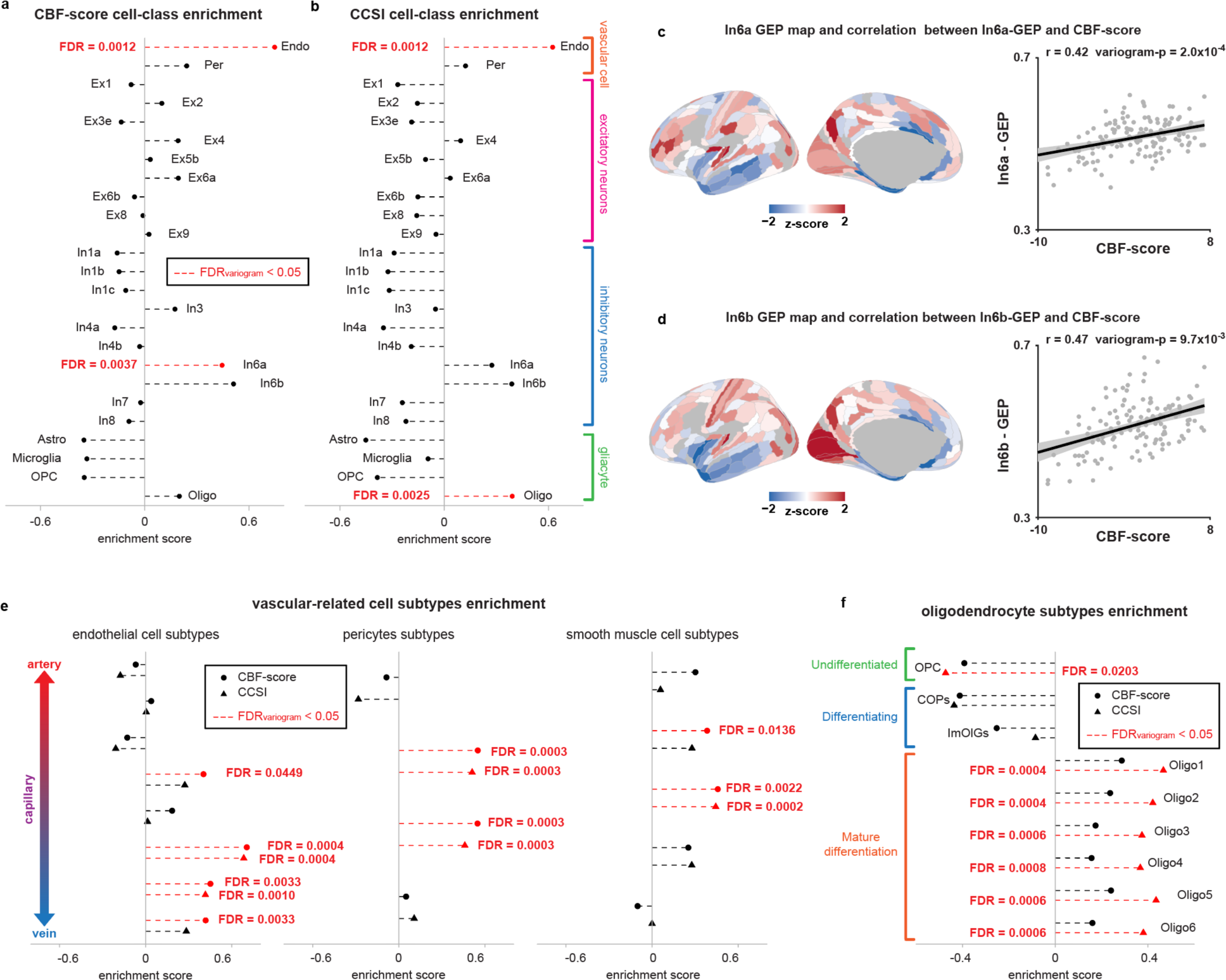
Cell-class enrichment analysis of CBF-score and CCSI by CGEM. Whole cell-class enrichment analysis of CBF-score (**a**) and CCSI (**b**) by CGEM. We used the variogram-null model to calculate the p value considering spatial autocorrelation. The exact FDR values that were significant after BH-FDR correction are marked in red (likewise in subsequent panels). Astro, Astrocytes; Oligo, Oligodendrocytes; OPC, Oligodendrocyte precursor cells; Endo, Endothelial cells; Per, Pericytes; Exa, Excitatory neuron subtype a; Ina, Inhibitory neuron subtype a. **c**, Spatial distribution of in6a neuron estimated using gene expression mapping and its relationship with CBF-score. Exact r and p values calculated by the variogram-null model are marked in the figure. The shading is the 95% confidence interval and dots represent individual brain region (same with **d**). **d**, Spatial distribution of in6b neuron estimated using gene expression mapping and its relationship with CBF-score. **e**, Vascular-related subtype cell-class enrichment analysis of CBF-score and CCSI by CGEM. These include various subtypes of three types of vascular cells: endothelial, pericyte, and smooth muscle cells. The vascular cells at the top come from arteries, the ones at the bottom come from veins, and the y-axis position of the capillaries in the middle also reflects the distance of each capillary cell from the artery or vein. **f**, Oligodendrocyte’s subtype cell-class enrichment analysis of CBF-score and CCSI by CGEM. COPs, Committed oligodendrocyte precursors; ImOlGs, Immature oligodendrocytes.

**Extended Data Fig. 7.**
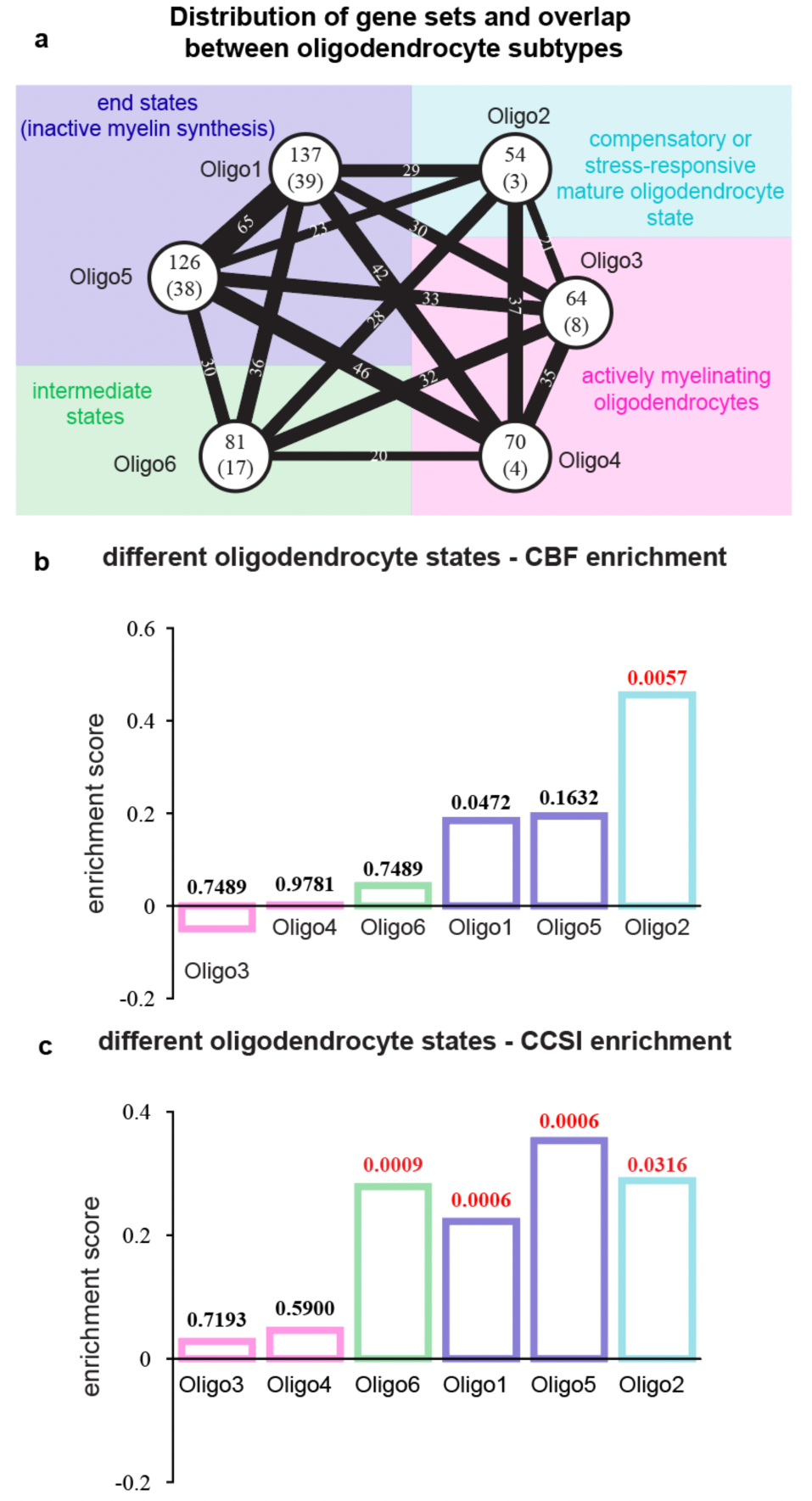
Different mature oligodendrocytes states enrichment analysis of CBF-score and CCSI. **a**, Distribution of gene sets and overlap between oligodendrocytes states. Each circle represents an oligodendrocyte in a certain state. The number in the circle indicates the number of genes in the marker gene set of that state. The number in the brackets indicates the number of genes remaining in the gene set after removing the overlap genes. The colored shading is based on the report of Jakel et al.^46^, which summarizes these six states into four functional states, including actively myelinating state, intermediate state, end state and oxidative stress state. **b**, Oligodendrocyte’s subtype cell-class enrichment analysis of CBF-score by redefined state-specific gene sets in **a**. The bar colors are consistent with the state classification in a. Exact FDR values calculated by the variogram-null model and BH-FDR correction are marked on each bar (same with **c**). **c**, Oligodendrocyte’s subtype cell-class enrichment analysis of CCSI by redefined state-specific gene sets in **a**.

**Extended Data Fig. 8.**
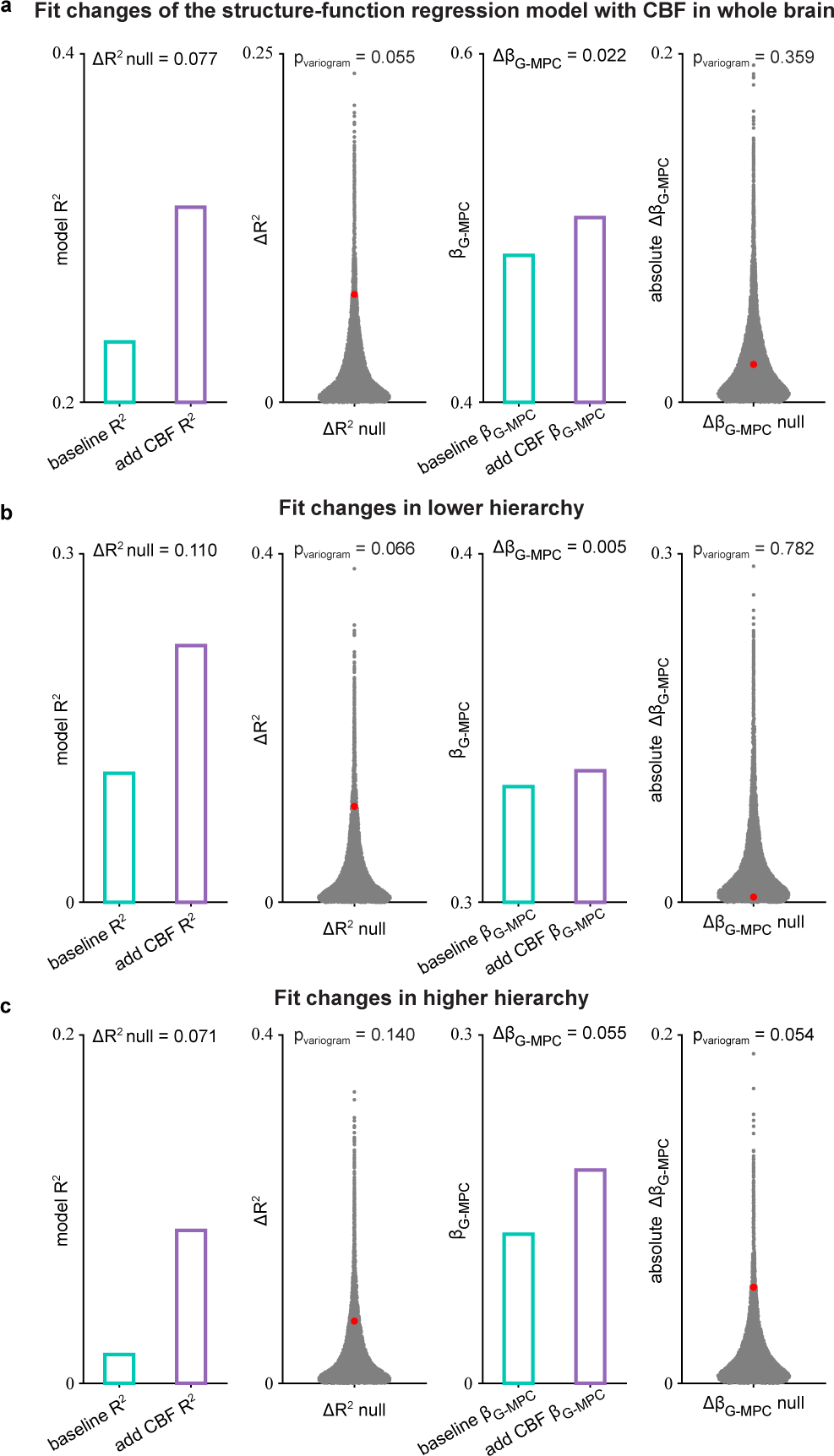
Fitting changes of the structure-function regression model with CBF. **a**, The fit changes of the structure-function regression model with CBF. Exact fitting parameters changes and p values calculated by the variogram-null model are marked in the figure. The cyan and purple bars represent the changes in the fitted parameters before and after CBF was added as a regression parameter. The gray dots represent the changes in the fitted parameters distribution calculated using the variogram-null model, and the red dots represent the positions of the true values in the null distribution (same with **b** and **c**). **b**, The fit changes of the structure-function regression model with CBF in lower hierarchy. **c**, The fit changes of the structure-function regression model with CBF in higher hierarchy.

